# Mobilization of nuclear antiviral factors by Exportin XPO1 via the actin network inhibits RNA virus replication

**DOI:** 10.1101/2024.12.20.629603

**Authors:** Biao Sun, Cheng-Yu Wu, Paulina Alatriste Gonzalez, Peter D. Nagy

**Author notes:** To whom correspondence and reprint requests should be addressed at the Department of Plant Pathology, University of Kentucky, 201F Plant Science Building, Lexington, KY 40546; /ext. 80726.

## Abstract

The intricate interplay between +RNA viruses and their hosts involves the exploitation of host resources to build virus-induced membranous replication organelles (VROs) in cytosol of infected cells. Previous genome- and proteome-wide approaches have identified numerous nuclear proteins, including restriction factors that affect replication of tomato bushy stunt virus (TBSV). However, it is currently unknown how cells mobilize nuclear antiviral proteins and how tombusviruses manipulate nuclear-cytoplasmic communication. The authors discovered that XPO1/CRM1 exportin plays a central role in TBSV replication in plants. Based on knockdown, chemical inhibition, transient expression and in vitro experiments, we show that XPO1 acts as a cellular restriction factor against TBSV. XPO1 is recruited by TBSV p33 replication protein into the cytosolic VROs via direct interaction. Blocking nucleocytoplasmic transport function of XPO1 inhibits delivery of several nuclear antiviral proteins into VROs resulting in dampened antiviral effects. The co-opted actin network is critical for XPO1 to deliver nuclear proteins to VROs for antiviral activities. We show that XPO1 and XPO1-delivered restriction factors accumulate in vir-condensates associated with membranous VROs. Altogether, the emerging theme on the role of vir-condensates is complex: we propose that vir-condensate serves as a central battleground between virus and the host for supremacy in controlling virus infection. It seems that the balance between co-opted pro-viral and antiviral factors within vir-condensates associated with membranous VROs could be a major determining factor of virus replication and host susceptibility. We conclude that XPO1 and nuclear antiviral cargos are key players in nuclear-cytoplasmic communication during cytosolic +RNA virus replication.

**Significance:** Tomato bushy stunt virus (TBSV), similar to other (+)RNA viruses, replicates in the cytosol and exploits organellar membrane surfaces to build viral replication organelles (VROs) that represent the sites of virus replication. The authors discovered that XPO1 exportin nuclear shuttle protein inhibited TBSV replication in plants. The conserved XPO1 is a central protein interaction nod, which propelled nucleocytoplasmic transport of several viral restriction factors into the cytosolic VROs that restricted tombusviruses replication. The delivered virus restriction factors provided inhibitory functions within virus-induced condensates associated with membranous VROs. The authors propose that the VRO-associated condensate serves as a central battleground between virus and the host for supremacy in controlling virus infection. Altogether, XPO1 is a critical protein interaction hub with major implications in viral replication. The authors conclude that XPO1 and its nuclear antiviral cargos are key players in nuclear-cytoplasmic communication during cytosolic (+)RNA virus replication.

## INTRODUCTION

Positive-strand RNA viruses are abundant and cause serious diseases as amply demonstrated by the recent emergence of flaviviruses and the SARS-CoV-2 global pandemic. +RNA viruses also cause major losses in agriculture and threaten global food security. (+)RNA viruses exploit the abundant resources of the host cells to build viral replication compartments/organelles (VROs) in the cytosol. The assembled VROs support +RNA virus replication in a membranous protective microenvironment [1–5].

Virus – host interactions are currently among the most intensively studied research areas due to the promising new antiviral approaches emerging from these studies [3, 5–10]. The host uses conserved innate and cell-intrinsic restriction factors as a first line of defense to fight viruses. Recent genome-wide screens with multiple viruses have identified dozens of restriction factors, which inhibit RNA virus accumulation [11–14]. Many restriction factors are nuclear RNA-binding proteins or shuttle proteins between the nucleus and the cytosol. How the nucleus might be able to mobilize nucleic acid binding or modifying proteins to fight against cytosolic RNA viruses is understudied.

Tomato bushy stunt virus (TBSV), a (+)RNA virus, is intensively studied to deepen our understanding of virus-host interactions, the mechanisms of virus replication and recombination [15–18]. An emerging theme with TBSV is that this virus dramatically remodels subcellular membranes and retargets various transport vesicles for VRO biogenesis [18–20]. Moreover, TBSV dramatically rewires cellular metabolism, and alters the lipid compositions of the targeted organelles [19–24]. TBSV co-opts the actin network and numerous host proteins, such as Hsp70 chaperone, translation factors eEF1A and eEF1Bγ, the ESCRT-associated proteins, Vps4 AAA ATPase and DEAD-box helicases needed for robust virus replication [25–31].

TBSV codes for two viral replication proteins, termed p33 and p92^pol^, which are essential for virus replication [32–34]. TBSV p33 protein with the help of co-opted host factors, rewires cellular pathways and optimizes the cellular milieu to support VRO biogenesis and viral RNA replication. Yet, the picture of virus-host interactions is further complicated by host responses, which include an arsenal of restriction factors to inhibit tombusvirus replication [16, 35–39].

Previous genome and proteome-wide studies on TBSV replication based on yeast model host identified exportin Crm1/XPO1, and other nuclear pore complex and export proteins, such as Nsp1, Nup1, Nup53, Nug1, Brl1, Ndc1, Snl1 and Sec13, and Srp40 transport chaperone and the importin Pse1 [16, 35–39]. It remains unclear if only the cytosolic pool of nuclear proteins plays a role in tombusvirus replication or if there is a mechanism(s) involving active protein translocation from the nucleus into the viral replication compartments. Identification of the nuclear shuttle protein, called exportin-1 (XPO1 or Crm1 in humans and yeast) suggested a putative role of nuclear export function in the replication of the cytosolic TBSV [35]. The evolutionarily conserved XPO1 is essential for exporting hundreds of proteins from the nucleus to the cytoplasm [40, 41]. This protein is so conserved in eukaryotes that the yeast XPO1/Crm1 can be complemented by human XPO1 [40, 41]. Among the XPO1 cargos are many nuclear RNA-binding proteins and putative antiviral factors [20, 42]. Nuclear export of proteins takes place through the nuclear pore complex with the help of nuclear export receptors, such as XPO1/Crm1 [40, 41]. XPO1 binds the nuclear cargo, usually carrying a specific leucine-rich nuclear export signal (NES). The active export of cargos is stimulated by Ran-GTP that facilitates XPO1 to bundle cargos as transport complexes, followed by export to the cytosol. Then, GTP-hydrolysis stimulated by co-factors RanGAP and RanBP1 facilitates the disassembly of the transport complex, releasing the cargos into the cytosol, whereas XPO1 is recycled to the nucleus for new rounds of export [43, 44].

XPO1 is an important protein and over-expressed in many forms of cancer [45, 46]. XPO1 is also important for HIV RNA transport out of the nucleus with the help of HIV Rev protein [47, 48]. Other nuclear viruses or their proteins, such as influenza virus, Nipah virus, Rhinovirus and DNA viruses depend on XPO1 to exit out of the nucleus [49–52]. Dengue virus NS5 protein enters the nucleus and interacts with XPO1, which is critical for modulation of host antiviral responses [53, 54]. XPO1 facilitates the nuclear export of the RdRp protein of Turnip mosaic virus from the nucleus to the cytosol [55, 56]. Thus, the emerging picture is that XPO1 is hugely important in cancer and viral diseases in humans [49].

In this paper, we studied the role of XPO1 in TBSV replication in plants and *in vitro*. Based on knockdown, chemical inhibition or over-expression and *in vitro* experiments, we showed that XPO1 acts as a cellular restriction factor against TBSV replication. XPO1 was found to be recruited by the TBSV p33 replication protein into the cytosolic VROs. BiFC and co-purification experiments demonstrated interaction between XPO1 and the viral p33 replication protein. We also demonstrated the nucleocytoplasmic transport function of XPO1 is needed to deliver nuclear antiviral proteins, such as DRB4, AGO2, nucleolin and CenH3, into VROs to inhibit TBSV replication. The co-opted actin network was critical in XPO1 antiviral activities. In addition, we showed that XPO1 and the XPO1-delivered restriction factors accumulated in vir-condensates associated with the membranous VROs. Altogether, we propose that XPO1 and its cargos are targeted into VROs, ultimately acting antiviral roles.

## RESULTS

### XPO1 exportin restricts tombusvirus replication in plants

Crm1/XPO1 has been identified in yeast screens as a putative antiviral host factor [35]. Because Crm1/XPO1 is a major protein interaction hub protein, we further tested its role in TBSV replication. We knocked down XPO1 mRNA level based on a virus-induced gene silencing (VIGS) approach in *Nicotiana benthamiana* [57]. Because XPO1 knockdown resulted in smaller plants than the control (TRV-cGFP) plants after 12 days, we shortened the VIGS experiments to 7 days (Fig. 1A). Replication of the peroxisome-associated TBSV and CNV (the closely-related cucumber necrosis virus) and the mitochondria-associated carnation Italian ringspot virus (CIRV) genomic (g)RNAs was increased by ∼2-to-3-fold in XPO1 knockdown (KD) plants when compared to the nonsilenced control plants two-three days after inoculation (Fig. 1B-D). The symptoms caused by all three tombusviruses became more severe in XPO1 KD than in control plants (Fig. 1B-D). Thus, the XPO1 KD plants are highly supportive of tombusvirus replication, suggesting that XPO1 functions as a restriction factor for tombusvirus replication.

**Figure 1.**
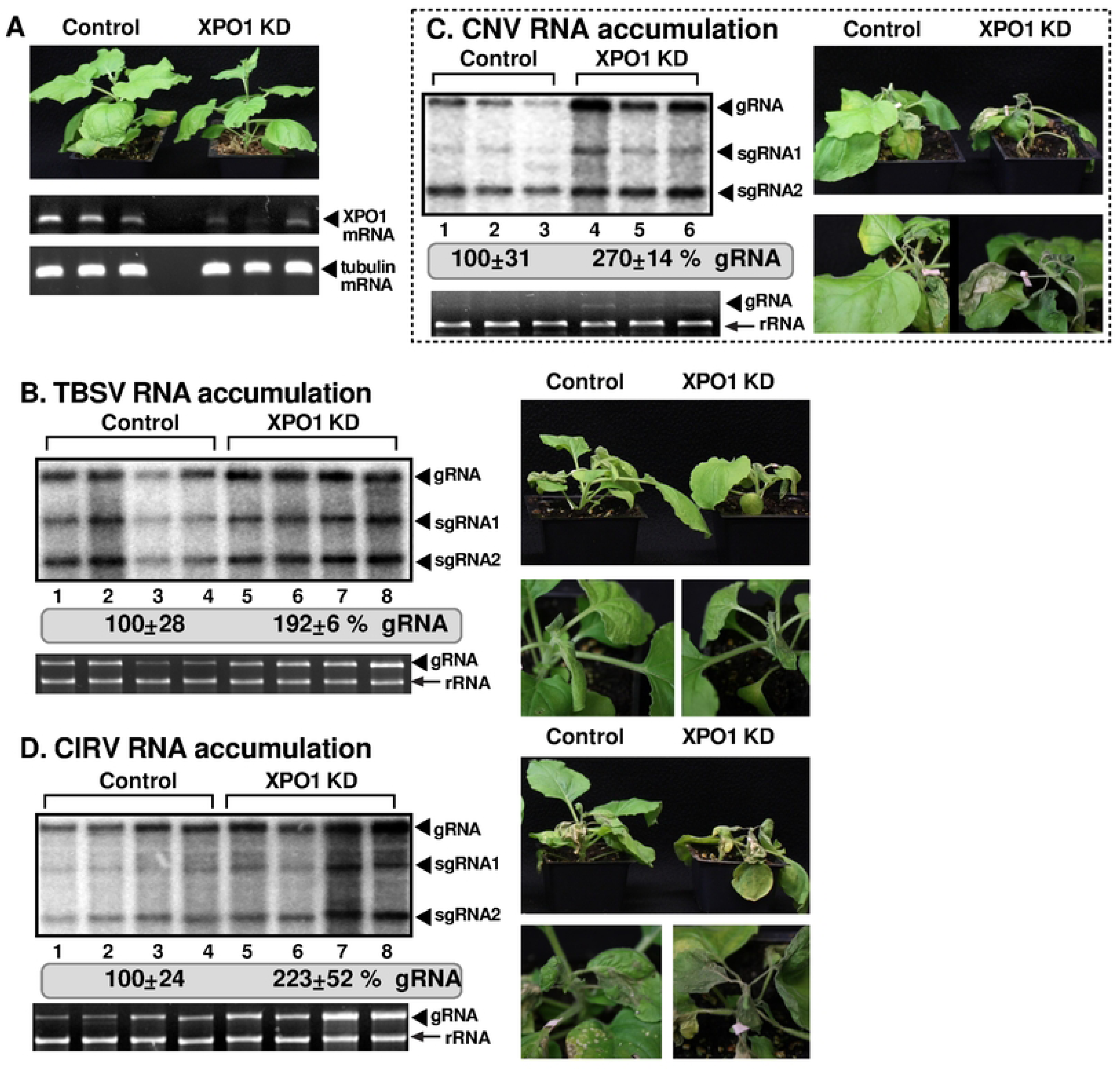
Knockdown of *XPO1* gene expression leads to enhanced tombusvirus replication in *N. benthamiana*. (A) Top panel: The phenotypes of *XPO1*-silenced *N. benthamiana* is shown at 7 days post agroinfiltration (7 dpai). VIGS was performed by agroinfiltration of tobacco rattle virus (TRV) vectors carrying partial sequence of *XPO1* or 3’ terminal GFP sequence as the control. Middle panel: RT-PCR analysis of XPO1 mRNA level in the silenced and control plants. Bottom panel: RT-PCR analysis of TUBULIN mRNA level in the silenced and control plants. (B) Accumulation of TBSV genomic (g)RNA and sgRNAs at 1.5 days after TBSV inoculation in XPO1-silenced (KD) plants was measured by northern blot analysis. The inoculation with TBSV was performed 7 days after VIGS. Second panel: The ribosomal RNA is shown as the loading control in agarose gel stained with ethidium-bromide. TBSV gRNA is visible in the gel. Right panel: Accelerated and more severe TBSV-induced symptom development was observed in XPO1-silenced *N*. *benthamiana.* The symptoms were documented at 5 days post TBSV inoculation. (C&D) Left panels: Accumulation of CNV or CIRV gRNA in XPO1-silenced *N. benthamiana* plants at 2 d post inoculation (dpi) was measured by northern blot analysis. See further details in panel B. Right panels: Accelerated and more severe CNV and CIRV-induced symptom development was observed in XPO1-silenced *N*. *benthamiana.* The pictures were captured at 5 dpi. Each experiment was repeated three times.

To further test the restriction function of XPO1 during tombusvirus replication in plants, we transiently expressed AtXPO1 in *N. benthamiana* via agroinfiltration. The same leaves were separately inoculated with the three tombusviruses 18 h later. RNA analysis showed ∼3-to-5-fold reduction in TBSV, CNV and CIRV gRNAs accumulation in the inoculated leaves (Fig 2A, C-D).

**Figure 2.**
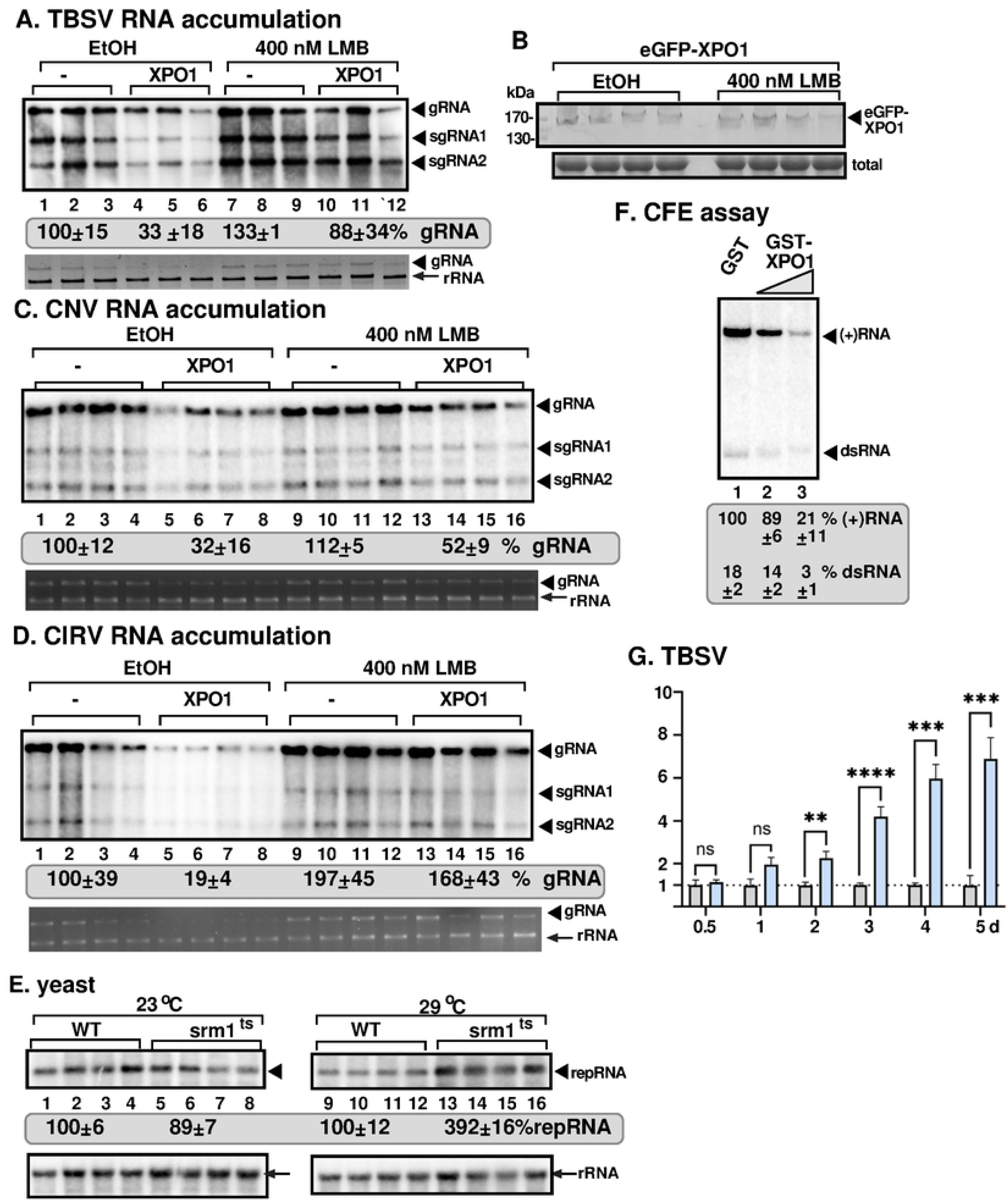
Expression of XPO1 inhibits tombusvirus accumulation in *N*. *benthamiana*. (A, C, and D). Transient expression of XPO1 in *N. benthamiana* inhibits accumulation of TBSV, CNV, and CIRV, respectively. Top panels: Accumulation of TBSV, CNV or CIRV RNAs at 1.5 dpi, 2 dpi and 2 dpi, respectively, was measured by northern blot analysis. Bottom panel: The 18S ribosomal RNA is shown in an agarose gel stained with ethidium-bromide as the loading control. *N. benthamiana* leaves were agroinfiltrated for transient expression of XPO1 (pGD vector as control), in combination with with 0.5% ethanol (EtOH) as a control or 400 nM of Leptomycin B (LMB), a chemical inhibitor of XPO1. The LMB and EtOH treatments were repeated 24 h later. (B) Expression of XPO1 was measured by western blot using anti-GFP antibody. Coomassie Brilliant Blue staining was used for the normalization of total proteins as loading control. (E) Accumulation of TBSV replicon repRNA in *srm1^ts^* or wild type (wt) yeasts at permissive temperature (23 ℃) or semi-permissive temperature (29 ℃) was measured with northern blotting. His_6_-p33 and His_6_-p92 RdRp were expressed from CUP1 promoter and TBSV (+)repRNA from GAL1 promoter to launch replication of TBSV repRNA. Top panel: Northern blot analysis of TBSV repRNA accumulation 1.5 d time point was done using a 3’ end specific probe. Bottom panel: The 18S ribosomal RNA is shown as the loading control, which was detected by northern blot. (F) Addition of XPO1 inhibits *in vitro* replication of TBSV repRNA based on TBSV replicase reconstitution assay in yeast cell-free extract (CFE). The *E. coli*-expressed and affinity-purified recombinant TBSV p33 and p92^pol^ replication proteins and TBSV (+)repRNA template were added to program CFE to support *in vitro* replication of TBSV repRNA. Increasing amounts (1.9 and 3.8 μM) of GST(as a control) or GST-XPO1 were added to the reactions. 5% polyacrylamide PAGE containing 8 M urea shows the produced ^32^P-labeled TBSV (+)repRNAs and dsRNA replication intermediate. (G) The transcriptional level of *XPO1* is induced during TBSV replication. Transcriptional levels of *XPO1* at six time points were estimated by RT-qPCR in total RNA samples obtained from mock-treated or TBSV-inoculated *N. benthamiana*. Each experiment was repeated three times.

Nucleus to cytosol export function of XPO1 can be blocked by inhibitors [58]. We treated the tombusvirus-inoculated leaves with Leptomycin B (LMB), which is a highly efficient and selective inhibitor of nuclear export. LMB binds to the NES-binding groove of XPO1 [40, 58]. Interestingly, replication of TBSV and CIRV was enhanced by ∼1.5-to-2-fold after LMB treatment of plant leaves (Fig. 2A&D). LMB treatment also neutralized the restriction function of XPO1 expressed transiently in the treated and TBSV, CNV or CIRV inoculated leaves (Fig. 2A, C&D). The above data demonstrate that the restriction function of the plant XPO1 depends on the nucleocytoplasmic export function in tombusvirus replication. The enhanced replication of TBSV and CIRV after the LMB treatment indicates that XPO1 restriction function might be due to the nucleocytoplasmic export of one or more restriction factors from the nucleus to the cytosol, where they might inhibit tombusvirus replication.

The nucleocytoplasmic transport function of XPO1 also depends on regulatory factors in the nucleus. These factors include Ran GEF (guanine nucleotide exchange factor, called Srm1/Prp20 in yeast and RCC1 in humans) that promotes exchange of GDP to GTP in RanGTPase. Then, RanGTP facilitates the formation of XPO1-RanGTP-cargo complex prior to transportation out of the nucleus through the nuclear pore [43, 44]. To test if this activity of RanGTP and XPO1 affects TBSV replication, we used a temperature-sensitive (ts) mutant of Srm1 in haploid yeast [59]. Growing the mutant srm1^ts^ yeast at the semi-permissive temperature (29 °C) resulted in ∼4-fold increased TBSV replication in comparison with yeast grown at the permissive temperature (Fig. 2E). Thus, the RanGTP function and nucleocytosolic transport seems to be important for XPO1 to operate as a tombusvirus restriction factor in yeast.

To further test the restriction function of XPO1, we reconstituted TBSV replicase *in vitro* using yeast cell-free extract (CFE) programmed with purified p33, p92^pol^ and a replicon (+)RNA [30, 60]. In the CFE assay, we used purified recombinant XPO1, which was added in increasing amounts at the beginning of the assay. At the end of the assay, we performed nondenaturing PAGE analysis of the *in vitro* replicase products. The replication assay revealed up to ∼5-fold reduction in both dsRNA replication intermediate [61] and in (+)ssRNA products in CFE with the highest amount of XPO1 in comparison with the RNA replication supported by WT CFE in the presence of GST control (Fig. 2F, lanes 2-3 versus 1). Inhibition of both (-)RNA synthesis (i.e., dsRNA production) and (+)RNA synthesis by XPO1 suggests that XPO1 blocks the TBSV replicase assembly steps, which occur prior to (-)RNA and (+)RNA synthesis *in vitro* [60, 62]. Because the CFE preparations are mostly free of the nuclear fraction, and XPO1 is purified from *E. coli*, it is likely that XPO1 has a direct restriction function against TBSV.

We found that transcription of XPO1 mRNA level was slowly increased during TBSV replication up to 7-fold by the fifth day of infection (Fig. 2G). This might indicate that *N. benthamiana* increases nuclear protein export during TBSV replication as an antiviral response.

### XPO1 interacts with the tombusvirus replication proteins and is recruited into the tombusvirus replication organelles in plants

TBSV replicates in the cytosolic side of clustered peroxisomes by assembling large viral replication organelles (VROs) [63–66]. To test if the restriction function of XPO1, which is a shuttle protein between the nucleus and cytosol (Fig. 3C) [43], is performed in the cytosol, we co-expressed TBSV p33-BFP replication protein and the eGFP-tagged XPO1 in *N. benthamiana* leaves. Confocal laser microscopy analysis revealed the co-localization of p33-BFP and eGFP-XPO1 within the cytosolic VROs (marked by the peroxisomal marker, RFP-SKL) in *N. benthamiana* cells replicating TBSV (Fig. 3A). We also used transgenic *N. benthamiana* expressing the H2B-CFP (Histon2B) nuclear marker protein, which showed that eGFP-XPO1 partitioned between the nucleus and VROs in plant cells infected with TBSV (Fig. 3B). Comparable studies with the CIRV p36 replication protein also showed the partial re-localization of eGFP-XPO1 to the CIRV VROs marked by CIRV p36-BFP or RFP-CoxIV mitochondrial marker protein (Fig. 3D-E). Distribution of eGFP-XPO1 in a single cell showed partitioning of XPO1 between the mitochondria-associated VROs and the nucleus in CIRV-infected cells (Fig. 3F). Based on these results, we suggest that tombusvirus infections of *N. benthamiana* plants induce the partial re-localization of XPO1 nuclear shuttle protein into the cytosolic VROs.

**Figure 3.**
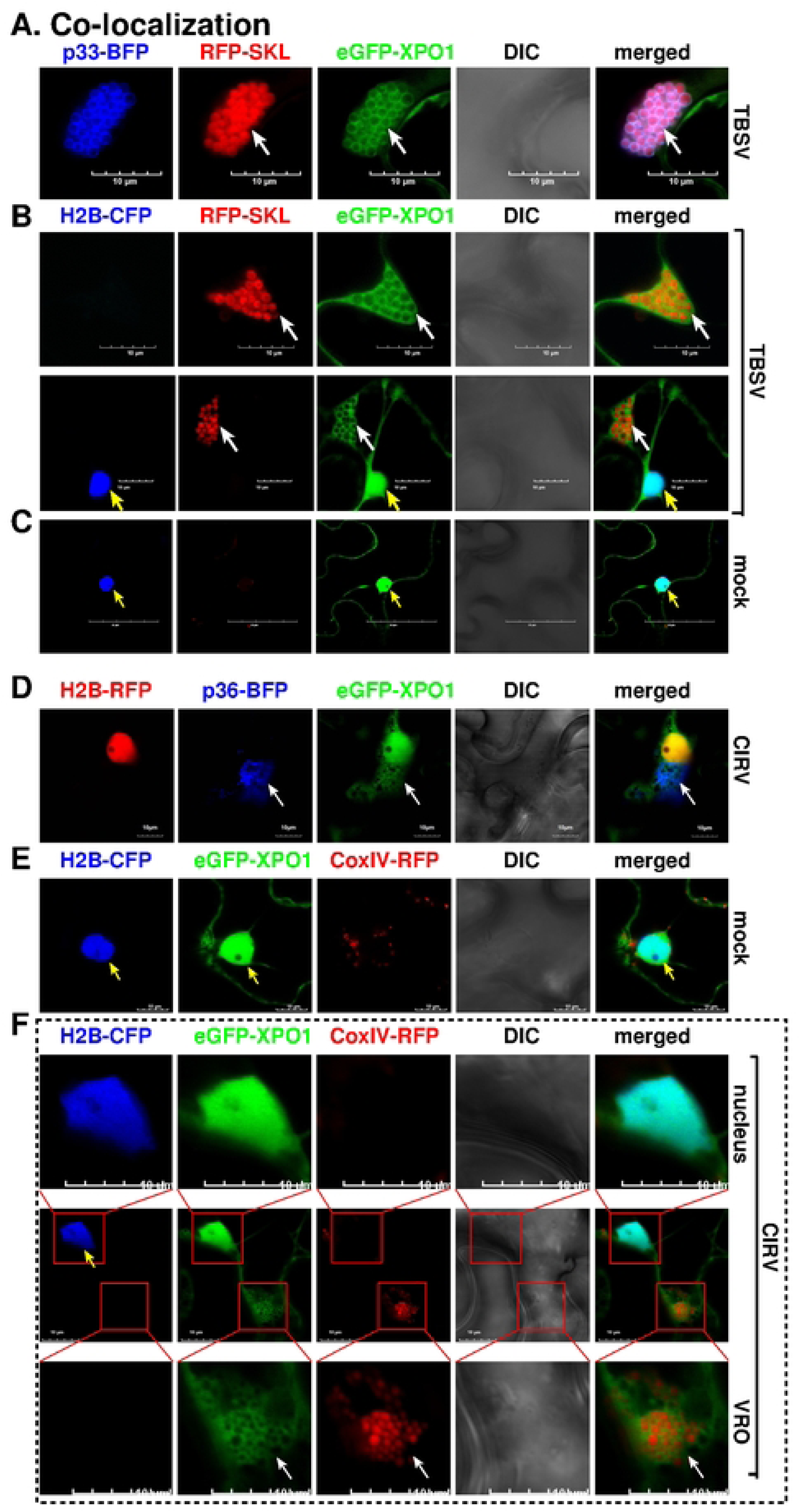
Recruitment of XPO1 by tombusviral replication protein into VROs in *N. benthamiana*. (A) Confocal laser microscopy images show co-localization of TBSV p33-BFP replication protein and eGFP-XPO1 in *N. benthamiana* during TBSV replication. The VROs consisting of clustered peroxisomes are indicated by a RFP-SKL peroxisomal marker. Scale bars represent 10 μm. (B) eGFP-XPO1 partitions between the nucleus and cytosolic VROs in plant cells during TBSV replication. Transgenic *N. benthamiana* expressing H2B-CFP (Histone2B) was used to mark the nucleus. VROs were marked by white arrows, whereas nucleus was pointed at by yellow arrows. Scale bars represent 10 μm. (C) eGFP-XPO1 distributes in the nucleus and cytosol in healthy transgenic H2B-CFP *N. benthamiana*. Scale bars represent 50 μm. (D) Confocal laser microscopy images show co-localization of CIRV p36-BFP replication protein and eGFP-XPO1. Transgenic *N. benthamiana* expressing H2B-RFP as the nuclear marker was used. Scale bars represent 10 μm. (E) The subcellular localization of eGFP-XPO1 in the mock-treated *N. benthamiana*. Mitochondria are indicated by CoxIV-RFP. Scale bars represent 10 μm. (F) Note that eGFP-XPO1 partitioned between the nucleus and VROs during CIRV replication. Middle panel: The nuclear localization and relocation to VROs of eGFP-XPO1 were captured in the same plant cell. Top panel: Enlarged images of the nuclear area. Bottom panel: Enlarged images of the VRO region. Scale bars represent 10 μm. Each experiment was repeated three times.

To test if the re-localization of XPO1 to VROs is dependent on the TBSV p33 replication protein, we used a BiFC approach in *N. benthamiana* plants. BiFC signals were observed within TBSV VROs, which formed from aggregated peroxisomes and located either at perinuclear or cytosolic locations (Fig. 4A). Similarly, we observed BiFC signals induced by the CIRV p36 replication protein together with XPO1 at the mitochondrial VROs (Fig. 4C). The negative control showed no BiFC signal (Fig. 4B). We conclude that the interaction between tombusvirus replication proteins and XPO1 mostly takes place in VROs that are frequently close to perinuclear regions.

**Figure 4.**
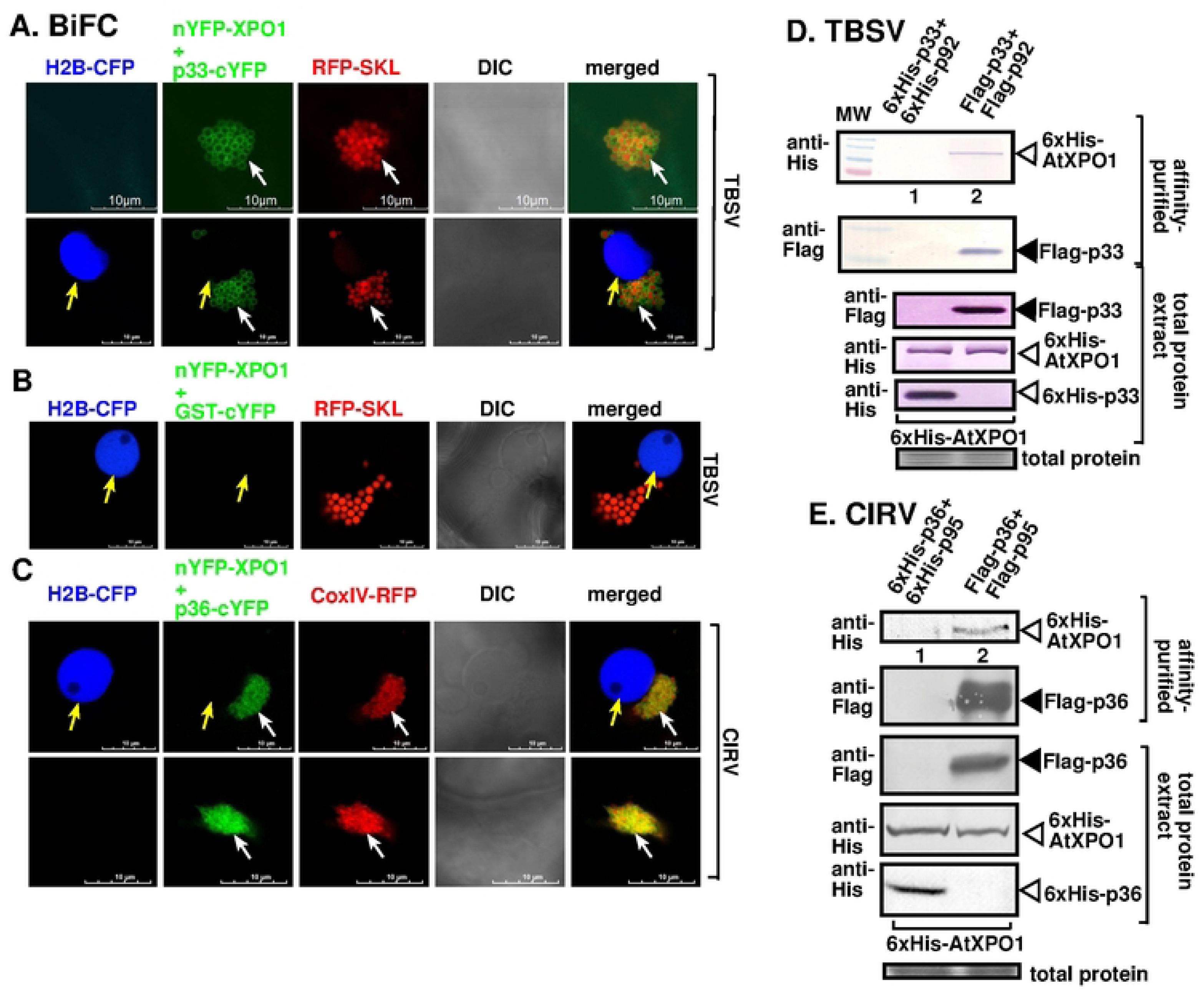
XPO1 interacts with tombusvirus replication proteins. (A) Interaction between TBSV p33-cYFP replication protein and nYFP-XPO1 was detected by BiFC during TBSV replication in H2B-CFP transgenic *N. benthamiana*. The green interaction signals surrounding the RFP-SKL marked peroxisomes are indicated by white arrows, demonstrating that interaction between p33-cYFP and nYFP-XPO1 occurs in VROs *in planta*. Nucleus is marked with yellow arrows. Scale bars represent 10 μm. (B) Control BiFC experiments with the combination of GST-cYFP and nYFP-XPO1 was performed in H2B-CFP transgenic *N. benthamiana* infected with TBSV. (C) Interaction between CIRV p36-cYFP and nYFP-XPO1 was identified by BiFC during CIRV replication in H2B-CFP transgenic *N. benthamiana*. The VROs, consisting of clustered mitochondria, are visualized by CoxIV-RFP mitochondrial marker. The interaction signals in VROs are indicated by white arrows. (D) Copurification of TBSV Flag-p33 and Flag-p92^pol^ replication proteins with His_6_-XPO1 from subcellular membranes of yeast. Top two panels: Co-purified His_6_-XPO1 (lane 2) and Flag-affinity-purified Flag-p33 were detected using western blot analysis. The negative control (lane 1) was based on yeast expressing His_6_-p33 and His_6_-p92^pol^ together with His_6_-XPO1. Middle three panels: Protein expression levels in total samples of yeast were detected by western blotting using the shown antibodies. Bottom panel: Coomassie Brilliant Blue staining was used for the normalization of total proteins as loading control. (E) Copurification of CIRV Flag-p36 and Flag-p95^pol^ replication proteins with His_6_-XPO1 from subcellular membranes of yeast. Top two panels: Co-purified His_6_-XPO1 (lane 2) and Flag-affinity-purified Flag-p36 were detected using western blot analysis. The negative control yeasts expressed His_6_-p36 and His_6_-p95^pol^ together with His_6_-XPO1. Middle three panels: Identification of protein expression in total samples from yeasts were detected by western blotting using the shown antibodies. See panel D. Each experiment was repeated three times.

To confirm direct interaction between XPO1 and either TBSV p33 or CIRV p36 replication proteins, we performed co-purification experiments from yeast co-expressing Flag-tagged p33, Flag-p92^pol^ or CIRV Flag-p36/Flag-p95^pol^ replication proteins and His_6_-tagged XPO1. After detergent-solubilization of the membrane-fraction of yeast, the Flag-tagged replication proteins were immobilized on a Flag-column. Western blot analysis of the eluted proteins from the column identified the co-purified His_6_-XPO1 (Fig. 4D-E, lane 2). These co-purification experiments demonstrated the interaction involving TBSV p33 or CIRV p36 replication proteins and XPO1 occurs in the yeast membrane fraction.

### Nucleocytoplasmic shuttling of XPO1 is needed for XPO1 re-localization to VROs

We blocked the cargo exporting function of XPO1 by applying LMB inhibitor, which specifically binds to XPO1, preventing the binding of cargoes in the nucleus [43]. Confocal microscopy of *N. benthamiana* plants infected with either TBSV or CIRV and expressing eGFP-XPO1 revealed poor re-localization of XPO1 from the nucleus into VROs in LMB-treated plant cells (Fig. 5A-B). This XPO1 distribution pattern is different from the localization in VROs in the control EtOH treated plant cells infected with TBSV or CIRV. We conclude that the nucleocytoplasmic shuttling activity of XPO1 is critical for its recruitment into tombusvirus VROs.

**Figure 5.**
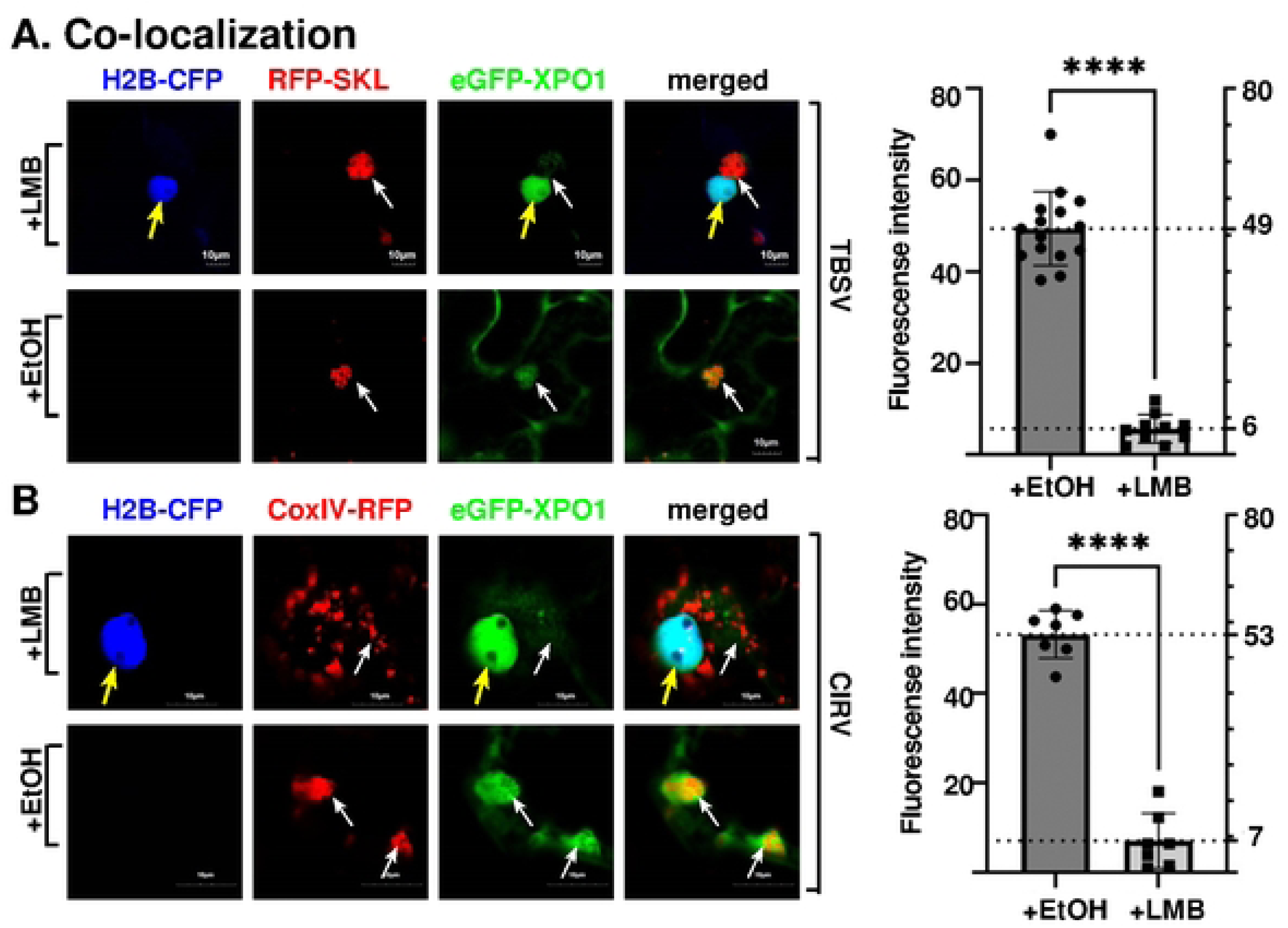
The relocation of XPO1 to cytosolic VROs depends on nucleocytoplasmic shuttling during tombusviruses replication. (A) Confocal microscopic images show the absence of relocation of eGFP-XPO1 to VROs during TBSV replication when treated with LMB inhibitor in H2B-CFP transgenic *N. benthamiana*. Bottom panel: Localization of eGFP-XPO1 when leaves were treated with 0.5% EtOH (control) in H2B-CFP transgenic *N. benthamiana*. Yellow arrows indicate the nucleus, while VROs are marked with white arrows. Scale bars represent 10 μm. Right panel: The graphs show fluorescence intensity of eGFP-XPO1 in VROs during TBSV replication after LMB versus EtOH treatments of H2B-CFP transgenic *N. benthamiana*. The fluorescent intensity in VRO regions is quantified by Olympus FV3000 FLUO-view software. T-test is used for data analysis utilizing GraphPad Prism 9 (**** represents P<0.0001). Error bars represent standard deviation (SD). (B) Confocal microscopic images show the absence of relocation of eGFP-XPO1 to VROs during CIRV replication when leaves were treated with LMB inhibitor in H2B-CFP transgenic *N. benthamiana*. The VROs, consisting of clustered mitochondria, are visualized by CoxIV-RFP mitochondrial marker. See further details in panel A. Each experiment was repeated three times.

### XPO1 delivers RNAi restriction factors from the nucleus to VROs during tombusvirus infections

Increased tombusvirus replication in XPO1 KD plants (Fig. 1) and by blocking of XPO1 re-location from the nucleus to the VROs by treatment with LMB (Fig. 2&5) indicate that a plausible explanation of anti-tombusvirus activity of XPO1 is the delivery of a pool of restriction factors from the nucleus to VROs. To test this model, we decided to characterize the effect of XPO1 on the re-localization of known nuclear restriction factors to tombusvirus VROs.

First, we studied selected components of the RNAi machinery, which are known to present in both nucleus and the cytosol [67, 68]. DRB4, which is a double-strand-RNA binding protein, is known to be re-targeted into TBSV VROs in *Arabidopsis* [69]. We observed that DRB4 was mostly restricted to nucleus after LMB treatment (S1D Fig), suggesting that DRB4 localization is affected by nucleocytoplasmic transport function of XPO1. We also found that a fraction of DRB4 was re-targeted to VROs during CNV (S1A-B Fig), TBSV (S2A-B Fig) or CIRV infections (S3A-B Fig). Interestingly, knocking down XPO1 expression or inhibiting XPO1 activity by LMB treatment reduced the recruitment of DRB4 into CNV VROs by ∼2.5-fold (Fig. 6A-B). Similarly, XPO1 significantly affected the recruitment of DRB4 into TBSV (S2E Fig) or CIRV VROs (S3E Fig). DRB4 expression inhibited CNV replication in *N. benthamiana* plants by 2-fold, confirming DRB4 antiviral function (Fig. 6C, lanes 5-8 versus 1-4). However, treatment of *N. benthamiana* leaves with LMB or knocking down XPO1 level interfered with the inhibitory effect of DRB4 on CNV replication (Fig. 6C-D). Altogether, we suggest that the antiviral function of DRB4 and its re-localization to the tombusvirus VROs depends on XPO1 nucleocytoplasmic export function.

**Figure 6.**
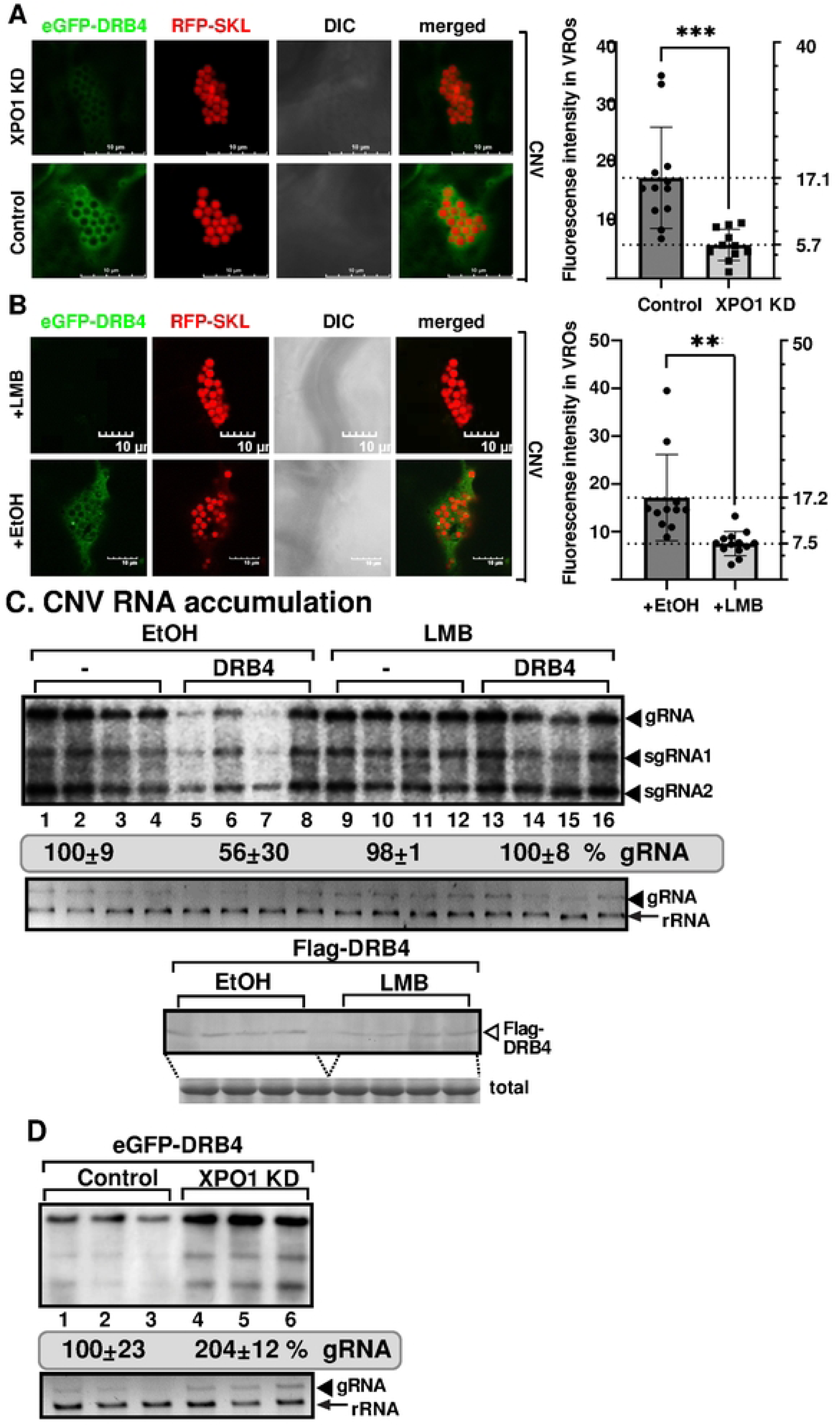
XPO1 delivers nuclear DRB4 to VROs to inhibit tombusvirus replication. (A) Recruitment of eGFP-DRB4 to VROs is reduced in XPO1 knockdown (KD) *N. benthamiana* infected with CNV. Left top panel: Confocal microscopic images show poor recruitment of eGFP-DRB4 into VROs (indicated by RFP-SKL) in XPO1 KD *N. benthamiana.* Bottom panel: Control experiment with *N. benthamiana* plants agroinfiltrated with the TRV vectors carrying partial GST sequence. Scale bars represent 10 μm. Right panel: Quantification of fluorescent intensity of eGFP-DRB4 in VRO regions was done with Olympus FV3000 FLUO-view software. T-test is used for data analysis utilizing GraphPad Prism 9 (*** represents P<0.001). Error bars represent standard deviation (SD). (B) Recruitment of eGFP-DRB4 to VROs is reduced in *N. benthamiana* infected with CNV and treated with LMB versus EtOH. See further details in panel A. (** represents P<0.01). (C) Transient expression of Flag-DRB4 in *N. benthamiana* inhibits CNV accumulation. Top panel: CNV RNA accumulation at 2 dpi was measured by northern blot analysis. *N. benthamiana* leaves were agroinfiltrated to express Flag-DRB4 (pGD vector as control) and the agroinfiltrated leaves were either infiltrated with 0.5% ethanol (EtOH) as a control or 400 nM of Leptomycin B (LMB). Middle panel: The 18S ribosomal RNA is shown in an agarose gel stained with ethidium-bromide as the loading control. Bottom panel: The expression of Flag-DRB4 was measured by western blot from the above samples using anti-flag antibody. Coomassie Brilliant Blue staining was used for the normalization of total proteins as loading control. (D) Top panel: Accumulation of CNV gRNA and sgRNAs at 2 dpi was measured by northern blot in *XPO1*-silenced plants expressing eGFP-DRB4. Agrobacteria infiltration to express eGFP-DRB4 (in the absence of p19 silencing suppressor) was performed 7 days after silencing of *XPO1* in *N. benthamiana*, followed two days later by inoculation with CNV. Bottom panel: The ribosomal RNA is shown as the loading control in the agarose gel stained with ethidium-bromide. Each experiment was repeated three times.

The second component of the RNAi machinery tested was AGO2. AGO2 protein is a component of the antiviral RISC complex, which targets plant RNA viruses, including tombusviruses [70, 71]. We expressed eGFP-AGO2 in *N. benthamiana* plants infected with CNV (S4 Fig), TBSV (S5 Fig) or CIRV (S5 Fig). Confocal microscopy analysis showed the efficient re-localization of AGO2 and XPO1 to VROs, representing either the clustered peroxisomes or mitochondria. As expected, AGO2 overexpression inhibited CNV replication (S7C Fig). However, treatment of *N. benthamiana* plants overexpressing AGO2 with LMB (S7C Fig) or knocking down XPO1 via VIGS (S7D Fig) dampened the antiviral activity of AGO2. LMB treatment or knocking down XPO1 expression significantly reduced the re-localization of AGO2 to CNV (S7A-B Fig), TBSV (S8A-B Fig) or CIRV VROs (S8C Fig). Based on these data, we conclude that the antiviral function and re-localization of AGO2 to tombusvirus VROs depends on XPO1 export function.

### XPO1 delivers cellular intrinsic restriction factors from the nucleus to VROs during viral infections

Tombusvirus replication is not only affected by the host powerful RNAi machinery, but cellular intrinsic restriction factors (CIRFs) limit tombusviruses in yeast and plants [14]. Several of the CIRFs are localized in the nucleus. Therefore, we tested if the recruitment of nuclear CIRFs is affected by XPO1 nucleocytoplasmic transport function. First, we studied the noncanonical function of CenH3 histone variant, which is recruited to tombusvirus VROs from the nucleus, resulting in inhibition of tombusvirus replication [72]. eGFP-CenH3 co-localized with RFP-XPO1 in TBSV (S9 Fig), CNV (S10 Fig) and CIRV VROs (S11 Fig) and nucleus. Treatment of *N. benthamiana* plants with LMB or knocking down XPO1 level inhibited the re-localization of CenH3 into TBSV (Fig. 7A-B) or CIRV VROs (S12 Fig) by ∼2.5-to-4-fold. Treatment of *N. benthamiana* leaves with LMB interfered with the inhibitory effect of CenH3 on TBSV (Fig. 7C) and CIRV replication (S12C Fig). Altogether, these data support the notion that the antiviral function and re-localization of CenH3 to the tombusvirus VROs depends on XPO1 function.

**Figure 7.**
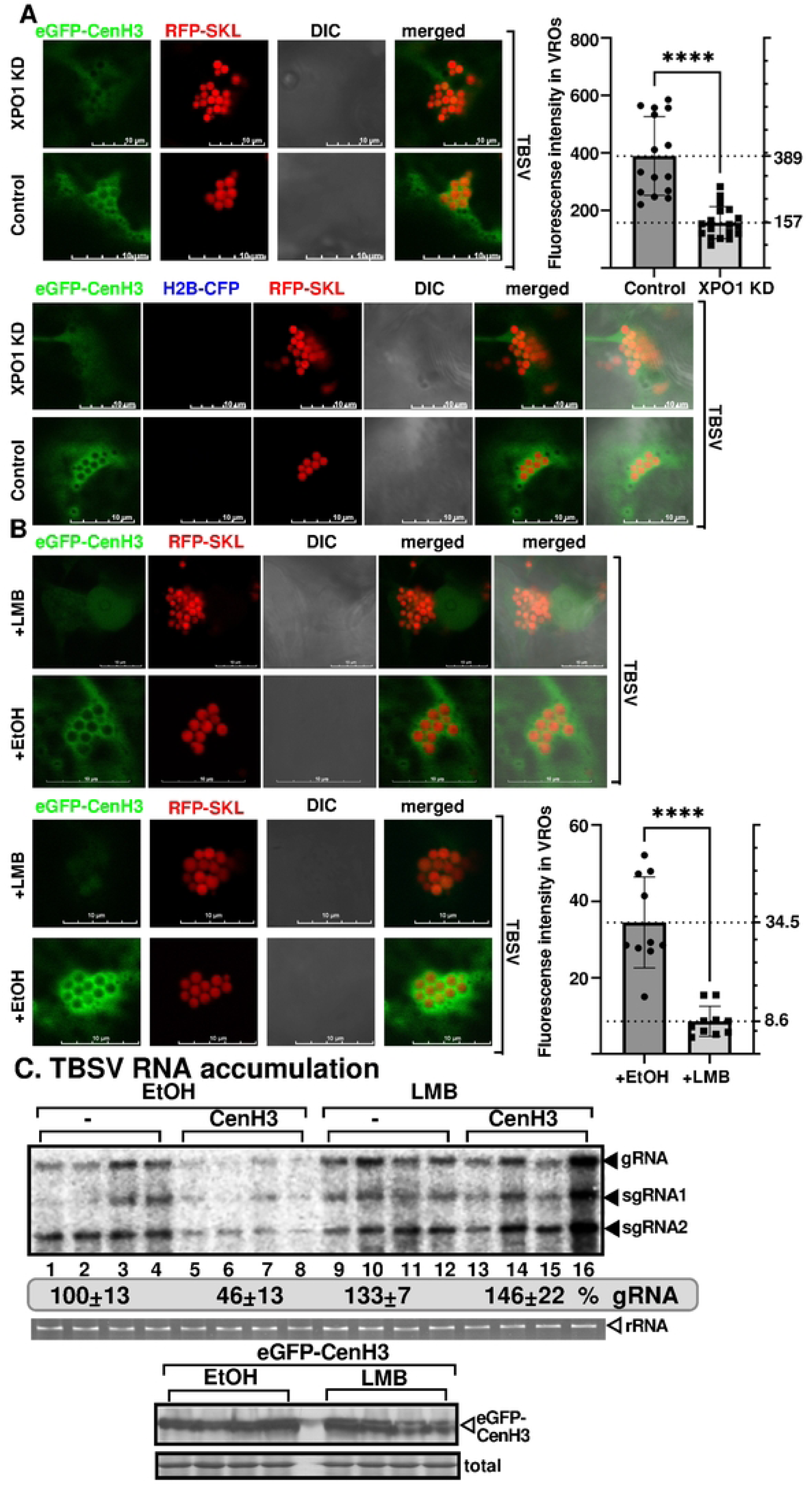
XPO1 delivers nuclear CenH3 to VROs to inhibit tombusvirus replication. (A) Recruitment of eGFP-CenH3 to VROs is reduced in XPO1 KD *N. benthamiana* infected with TBSV. Left top panel: Confocal microscopic images show poor recruitment of eGFP-CenH3 into VROs (indicated by RFP-SKL) in XPO1 KD *N. benthamiana.* Bottom panel: Control experiment with *N. benthamiana* plants agroinfiltrated with the TRV vector carrying partial GST sequence. Scale bars represent 10 μm. Right panel: Quantification of fluorescent intensity of eGFP-CenH3 in VRO regions with Olympus FV3000 FLUO-view software. See further details in Fig. 6A. (B) Recruitment of eGFP-CenH3 to VROs is reduced in *N. benthamiana* infected with TBSV and treated with LMB versus EtOH. Note that we show two independent sets of images to illustrate similar trends in different plants. See further details in Fig. 6A. (C) Transient expression of Flag-CenH3 in *N. benthamiana* does not inhibit TBSV accumulation in plants treated with LMB. Top panel: TBSV RNA accumulation at 1.5 dpi was measured by northern blot analysis. *N. benthamiana* leaves were agroinfiltrated to express Flag-CenH3 (or pGD vector as control) and the agroinfiltrated leaves were either infiltrated with 0.5% ethanol (EtOH) as a control or 400 nM LMB. Middle panel: The 18S ribosomal RNA is shown in an agarose gel stained with ethidium-bromide as the loading control. Bottom panel: The expression of Flag-CenH3 was measured by western blot from the above samples using anti-flag antibody. Coomassie Brilliant Blue staining was used for the normalization of total proteins as loading control. Each experiment was repeated.

The second nuclear CIRF tested was Nuc-L1 nucleolin (Nsr1 in yeast) that was previously shown to inhibit TBSV replication in yeast [73]. Nuc-L1 interacts with both p33 replication protein (S13B Fig) and the TBSV RNA [73]. Nuc-L1 also interacts with XPO1 mostly in the nucleus in *N. benthamiana* (S13A Fig). Interestingly, LMB treatment strongly inhibited the BiFC signal between Nuc-L1 and XPO1 in the nucleus (S13A Fig, bottom panels). Nuc-L1 interacted with XPO1 in both the nucleus and TBSV VROs in TBSV-infected control plants (S13A Fig, top panel). However, LMB-treatment strongly inhibited Nuc-L1 - XPO1 interaction in TBSV VROs (S13A Fig, second panel). LMB-treatment neutralized the inhibitory effect of Nuc-L1 over-expression on CNV accumulation in *N. benthamiana* (S13C Fig). Western-blotting showed that the LMB-treatment did not inhibit the expression of Nuc-L1 (S13C Fig). These data support the role of XPO1 in delivering Nuc-L1 antiviral restriction factor into tombusvirus VROs. Based on the above data, we conclude that several host restriction factors are delivered from the nucleus into tombusvirus VROs in an XPO1-dependent fashion.

### Critical role of the actin network in delivering XPO1 and antiviral cargos into tombusvirus VROs

The above data convincingly showed that XPO1 interacts with the tombusvirus replication proteins and delivers antiviral factors from the nucleus to VROs to restrict tombusvirus replication. How are XPO1 and the nuclear restriction factors targeted to VROs? One of the possible mechanisms is the contribution of the actin network. TBSV is known to hijack actin filaments to recruit cytosolic factors with pro-viral functions into VRO-associated vir-condensates [74–76]. TBSV p33 replication protein inhibits actin depolymerization function of Cof1/ADF1 actin filament disassembly protein, thus resulting in stable actin filaments and actin cables in TBSV-infected cells [63]. Therefore, we decided to test if the co-opted stabilized actin filaments could assist the movement of XPO1 and its cargos into VROs. Confocal imaging revealed the association of XPO1 with actin filaments within VROs and in the cytosol of TBSV- or CIRV-infected cells (Fig. 8A-B). Moreover, BiFC experiments showed that TBSV p33 and CIRV p36 replication proteins in complex with XPO1 were associated with actin filaments (Fig. 8D-E). We also observed that XPO1 cargos, such as DRB4 (S14A-B Fig), AGO2 (S14D-F Fig) and CenH3 (S15A-B Fig) were associated with actin filaments within VROs and in the cytosol of TBSV- or CIRV-infected cells. These observations highlighted the possibility that the co-opted actin filaments are used to deliver XPO1 and its nuclear cargos into VROs.

**Figure 8.**
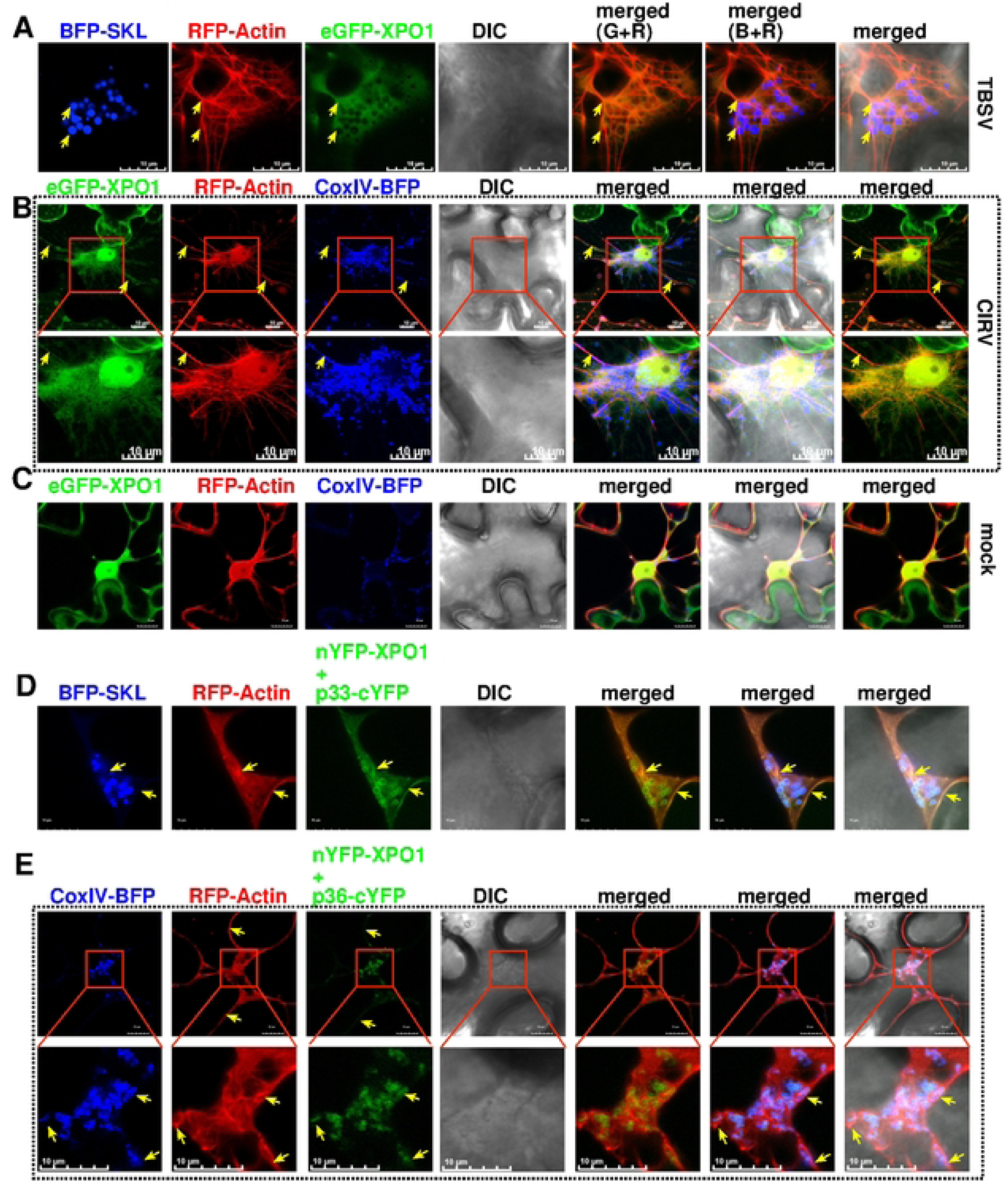
Association of XPO1 with coopted actin filaments during tombusvirus replication. (A-B) Confocal microscopy images show co-localization of eGFP-XPO1, RFP-Actin filaments and VROs, indicated by BFP-SKL during TBSV replication or by CoxIV-BFP mitochondrial marker during CIRV replication in *N. benthamiana*. The actin filaments were visualized by agro-expression of Lifeact, which binds to filamentous actin (F-actin) in plant cells. The actin filaments co-localized with eGFP-XPO1 are indicated by yellow arrows. The scale bar represents 10 μm. The enlarged images from the red boxed areas contain VROs and nuclear region. (C) Localization of eGFP-XPO1, RFP-Actin, and CoxIV-BFP mitochondrial marker in mock-treated *N. benthamiana*. The scale bar represents 10 μm. (D-E) Confocal microscopy images show the co-localization of RFP-Actin and BiFC interaction signals of nYFP-XPO1 and TBSV p33-cYFP or CIRV p36-cYFP in *N. benthamiana* cells. TBSV VROs are marked with SKL-BFP peroxisomal marker, whereas CIRV VROs are marked with CoxIV-BFP mitochondrial marker. The scale bar represents 10 μm. Each experiment was repeated.

To further test this model, we destroyed the cytosolic actin network by transiently expressing RavK effector of *Legionella* bacterium in *N. benthamiana* cells (Fig. 9C) [76–78]. Transient expression of RavK in cells infected with TBSV led to ∼2.5-fold reduced recruitment of XPO1 into TBSV VROs (Fig. 9A-B). Transient RavK expression also inhibited XPO1 recruitment into CIRV VROs formed by clustered mitochondria by ∼3-fold (S16A Fig). We also transiently expressed VipA effector of *Legionella* bacterium in *N. benthamiana* cells, which is an actin nucleator and stabilizes actin filaments (Fig. 9C) [79]. VipA expression is known to facilitate TBSV replication [76, 77]. Interestingly, transient expression of VipA in cells infected with TBSV led to ∼2-fold enhanced recruitment of XPO1 into TBSV VROs (Fig. 9A-B). These data suggest that recruitment of XPO1 into VROs is facilitated by co-opted actin filaments in TBSV-or CIRV-infected cells.

**Figure 9.**
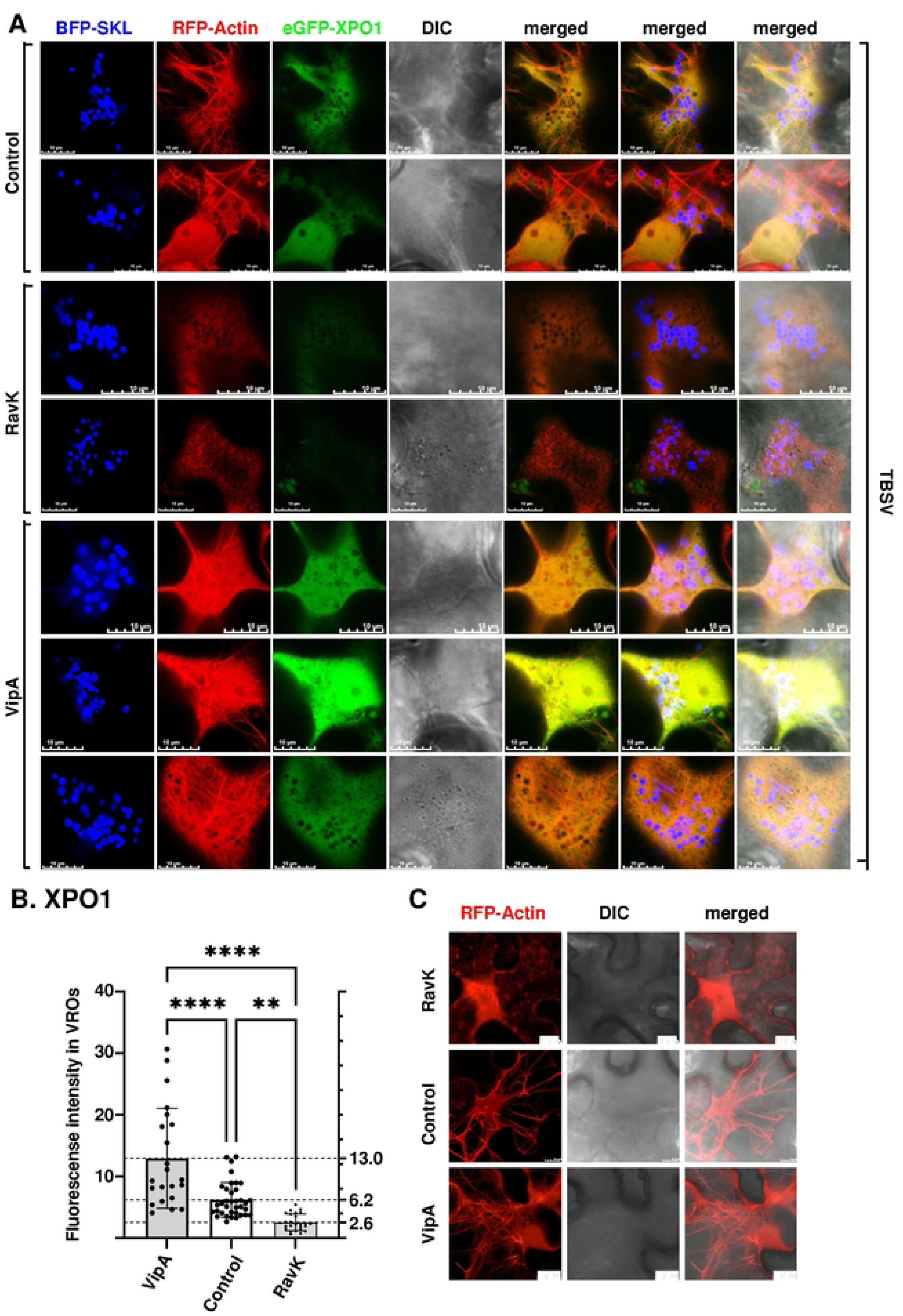
Recruitment of XPO1 to VROs depends on the co-opted actin filament networks. (A) Confocal microscopy images show recruitment of eGFP-XPO1 into VROs is affected by actin filaments during TBSV replication. Top panel: Co-localization of eGFP-XPO1, RFP-Actin filaments and VROs, indicated by BFP-SKL during TBSV replication in *N. benthamiana*. See further details in Fig. 8A. Middle two panels: Confocal microscopy images show the poor recruitment of eGFP-XPO1 into TBSV VROs when the actin filaments were destroyed by the transient expression of RavK effector of *Legionella* bacterium from a plasmid via agroinfiltration in *N. benthamiana*. Bottom three panels: Confocal microscopy images show efficient recruitment of eGFP-XPO1 into TBSV VROs when the actin filaments were induced and stabilized by the transient expression of VipA effector of *Legionella* bacterium from a plasmid via agroinfiltration in *N. benthamiana*. Scale bars represent 10 μm. (B) Quantification of eGFP-XPO1 within VROs based on experiments shown in panel A. The fluorescent intensity of eGFP-XPO1 in VRO regions is quantified by Olympus FV3000 FLUO-view software. T-test is used for data analysis utilizing GraphPad Prism 9 (**** represents P<0.0001, ** represents P<0.01). Error bars represent standard deviation (SD). (C) Control experiments to show distribution patterns of actin filament network when RavK or VipA effectors are transiently expressed versus no protein expression in *N. benthamiana.* Scale bars represent 10 μm. Each experiment was repeated.

Transient expression of RavK inhibited the delivery of XPO1 cargos DRB4 (Fig 10A-B), CenH3 (Fig 10C-D) and AGO2 (Fig 10 E-F) into TBSV VROs by ∼2-to-4-fold. Similarly, disruption of actin filaments by RavK inhibited the recruitment of DRB4 (S16C-D Fig), AGO2 (S17C-D Fig) and CenH3 (S17A-B Fig) into CIRV VROs by ∼2-to-4-fold. These data established that the co-opted actin network facilitates the delivery of XPO1 and its nuclear antiviral cargos into tombusvirus VROs to restrict tombusvirus replication in plants.

**Figure 10.**
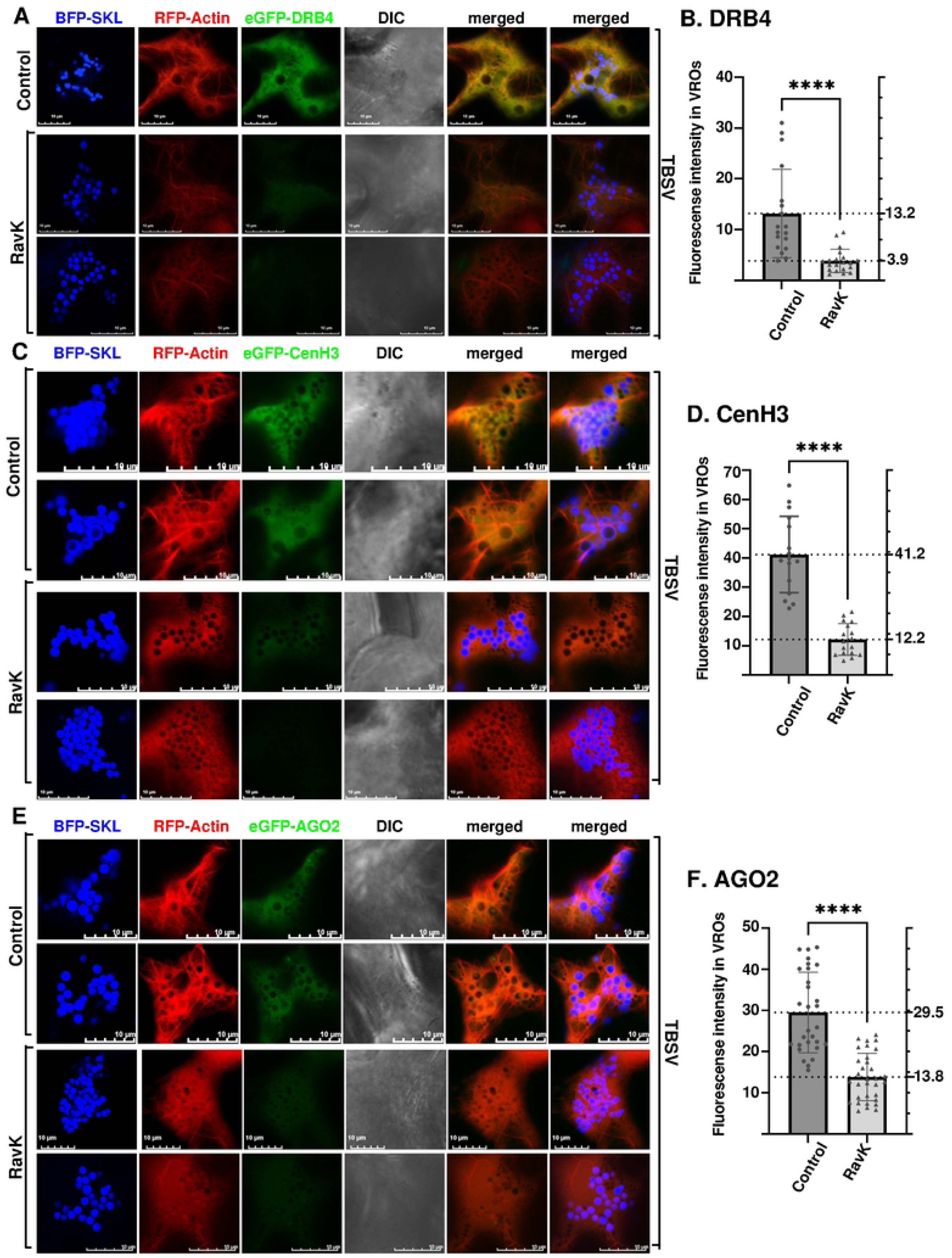
Coopted Actin filament network plays a critical role in delivering antiviral cargos of XPO1 to VROs. Confocal microscopy images show the poor recruitment of eGFP-DRB4 (A), eGFP-CenH3 (C), and eGFP-AGO2 (E) into TBSV VROs when the actin filaments were destroyed by the transient expression of RavK effector of *Legionella* bacterium from a plasmid via agroinfiltration in *N. benthamiana*. Top two panels in (A, C, and E) show the control confocal images without the expression of RavK effector in *N. benthamiana*. Scale bars represent 10 μm. Quantification of VRO-localized eGFP-DRB4 (B), eGFP-CenH3 (D) and eGFP-AGO2 (F) is shown in graphs. The fluorescent intensity in VROs is quantified by Olympus FV3000 FLUO-view software. T-test is used for data analysis utilizing GraphPad Prism 9 (**** represents P<0.0001). Error bars represent standard deviation (SD). Each experiment was repeated three times.

### Regulatory co-factors of XPO1 are recruited into tombusvirus VROs

The disassembly of the XPO1-RanGTP-cargo complex requires the conversion of RanGTP into a GDP-bound form (RanGDP) in the cytosol. This step is facilitated by RanGAP1 or RanGAP2 and RanBP1 factors [44, 80]. We found that RanGAP1/2 are recruited into TBSV (Fig. 11A-B) and CIRV VROs (S18A-B Fig). In addition, RanBP1 is also present in TBSV (Fig. 11C) and CIRV VROs (S18C Fig). Colocalization experiments showed that the recruitment of RanGAP1/2 (Fig 11D-E) and RanBP1 factors (Fig. 11F) into TBSV VROs takes place with the help of the actin network. Similarly, the actin network could play a role in the recruitment of RanGAP1/2 (S18D-E Fig) and RanBP1 factors (S18F Fig) to VROs consisting of aggregated mitochondria in CIRV-infected leaves. Recruitment of these cytosolic regulatory co-factors of XPO1 suggests that XPO1-RanGTP-cargo complex is likely disassembled in tombusvirus VROs, leading to the release of the antiviral cargos from the cargo complex in VROs. RanGAP1/2 and RanBP1 do not seem to interact with TBSV p33 or CIRV p36 replication proteins based on BiFC analysis in *N. benthamiana* (S19A-B Fig). These regulatory co-factors are likely recruited into VROs via interaction with XPO1.

**Figure 11.**
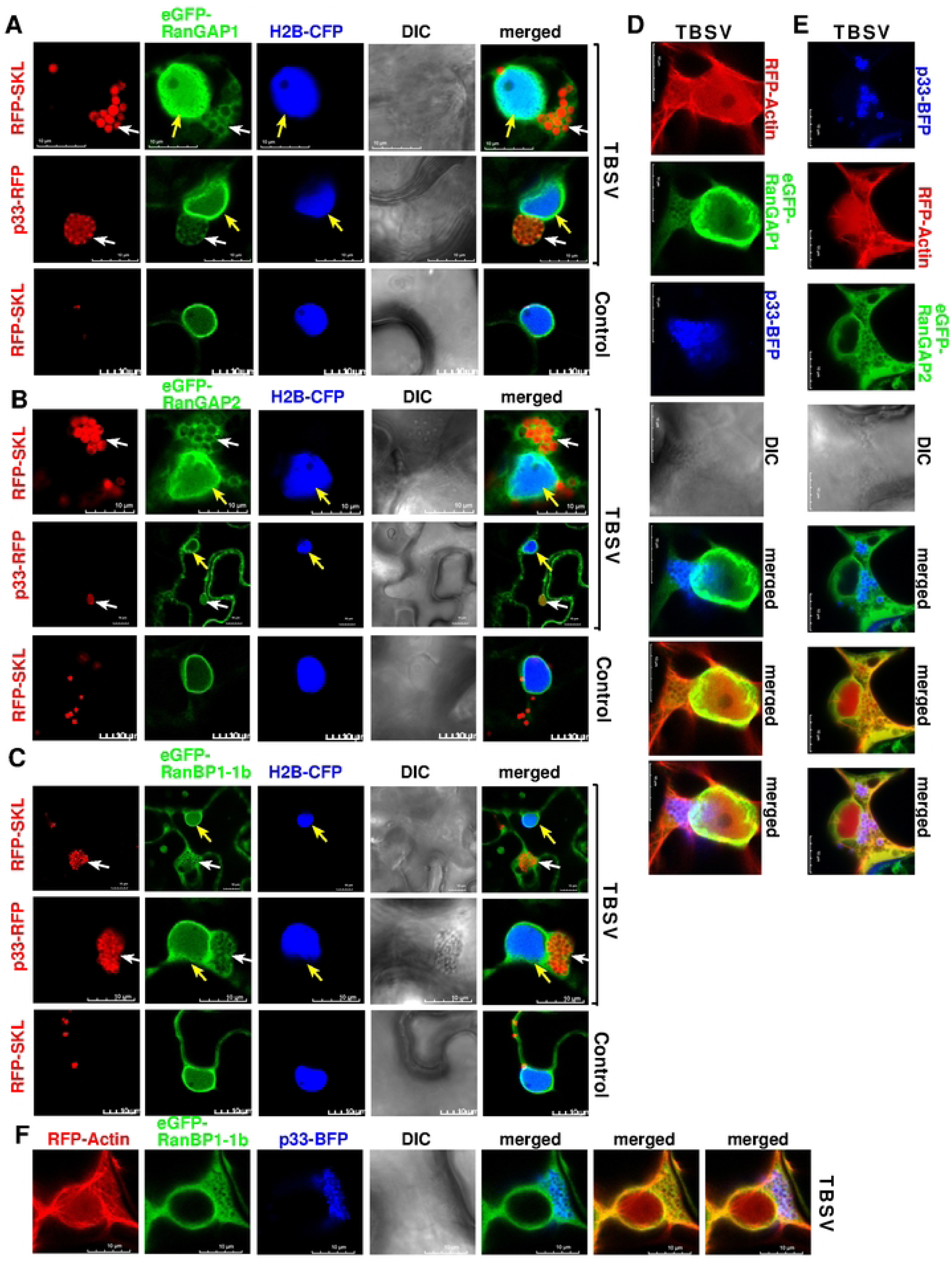
Regulatory co-factors of XPO1 are recruited into VROs during TBSV replication. Confocal microscopy images show the recruitment of eGFP-RanGAP1 (A), eGFP-RanGAP2 (B), or eGFP-RanBP1-1b (C) into VROs during TBSV replication. The bottom images in panels A, B and C show the subcellular localizations of eGFP-RanGAP1, eGFP-RanGAP2, or eGFP-RanBP1-1b in control plant cells. Yellow arrows indicate the nucleus, while VROs are marked with white arrows. Note these images were taken using H2B-CFP transgenic *N. benthamiana* marking the nucleus. Confocal microscopy images also show the associations of eGFP-RanGAP1 (D), eGFP-RanGAP2 (E), eGFP-RanBP1-1b (F) and p33-BFP replication protein with actin filament networks during TBSV replication. Scale bars represent 10 μm.

### Recruited XPO1 and antiviral factors are present in vir-condensates associated with membranous VROs

Tombusvirus VROs contain co-opted peroxisomal or mitochondrial membranes harboring the viral spherules in association with vir-condensates formed via p33-induced liquid-liquid phase separation of sequestered cytosolic proteins [74]. To test if XPO1 and the delivered antiviral cargos are present in vir-condensates, we performed FRAP experiments in *N. benthamiana* cells replicating TBSV. The slow recovery of the fluorescent signals after photobleaching of VROs suggested that XPO1 (Fig 12A) and the delivered DRB4 (Fig 12B), CenH3 (Fig 12C) or AGO2 (Fig 12D) cargos are all present in TBSV and CIRV (S20 Fig) vir-condensates within VROs. These observations suggest that the vir-condensate is the location where these antiviral factors perform their inhibitory function on tombusvirus replication. Thus, the host cells exploit the tombusvirus-induced vir-condensates to concentrate antiviral proteins in VROs as a virus-defense mechanism to restrict tombusvirus replication.

**Figure 12.**
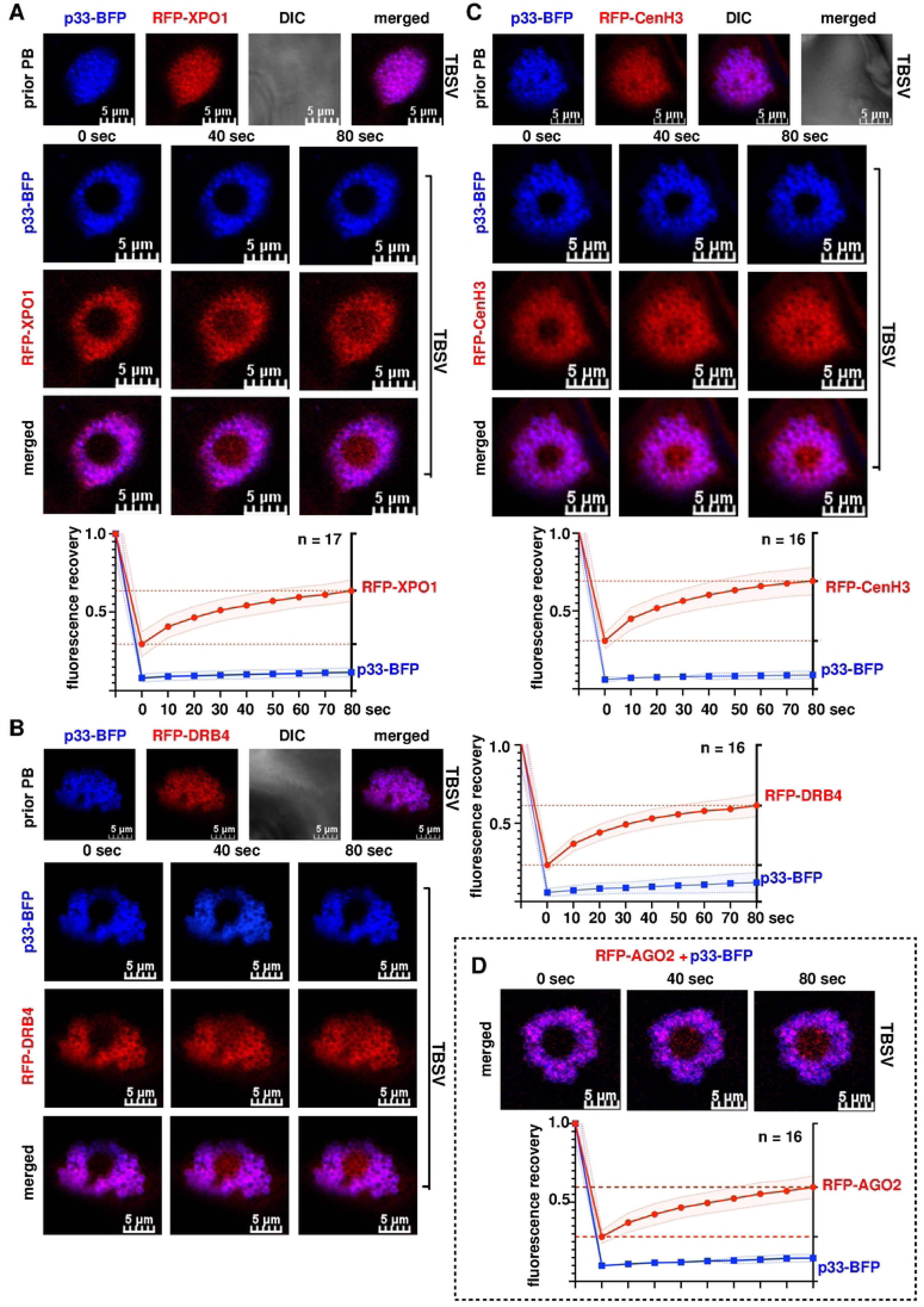
XPO1 and the antiviral cargos are present in vir-condensates associated with membranous VROs. (A) FRAP analysis shows partial fluorescence recovery of RFP-XPO1 after photobleaching in a single VRO in *N. benthamiana* during TBSV replication. Top panel: Co-localization of replication protein p33-BFP and RFP-XPO1 in VRO before photobleaching. Middle panels: Confocal microscopy images show partial fluorescence recovery of RFP-XPO1 at 0-, 40-, and 80-seconds post photobleaching during TBSV replication. Note that signals of p33-BFP did not recover in the FRAP assay because it is a membrane-bound protein. Scale bars represent 5 μm. Bottom panel: The graph shows time course analysis of FRAP data on RFP-XPO1 signal recovery in individual VROs during TBSV replication. Sample size *n* is annotated in the figure. Shaded area represents SD. Similar FRAP analysis of RFP-DRB4 (panel B), RFP-CenH3 (panel C), and RFP-AGO2 (panel D) are also shown. See further details in panel A. Each experiment was repeated three times.

## DISCUSSION

Tombusviruses, similar to other (+)RNA viruses, replicate in the cytosol and exploit organellar membrane surfaces to build VROs. However, previous genome- and proteome-wide studies of TBSV in the yeast model host, which supports TBSV replication, also identified numerous host factors localized mainly in the nucleus in the absence of viral components [32, 39, 81]. These findings indicated a yet unknown contribution of the nucleus to tombusvirus replication. Among the nuclear factors identified, XPO1 exportin shuttle protein was intriguing as a restriction factor. Mutation of XPO1/Crm1 in yeast enhanced TBSV replication, thus revealing a new layer in antiviral mechanisms [35]. But how could a nuclear shuttle protein affect the replication of a cytosolic virus?

XPO1 is a major protein interaction hub, which is involved in exporting ∼1,000 proteins and RNA-protein complexes out of the nucleus [40, 43]. Therefore, viral hijacking of XPO1 seems to provide easy access to protein rich resources, such as RNA-binding proteins and helicases, which are otherwise hidden in the nucleus from a cytosolic virus. However, among the cargo proteins of XPO1 could be restriction factors, which present a challenge to a simple (+)RNA virus with limited protein coding capacity, such as tombusviruses. Accordingly, data presented in this paper show that XPO1 acts as a restriction factor (CIRF) of tombusvirus replication for both the peroxisome-associated TBSV and CNV and the mitochondria-associated CIRV in *N. benthamiana* plants. The *in vitro* replicase reconstitution experiment showed that XPO1 directly inhibits TBSV replication, likely via binding and sequestering of p33 replication protein, thus preventing p33 to perform its essential functions during viral replication. However, *in planta* experiments with LMB inhibitor of XPO1 or knocking down XPO1 level via VIGS showed that the main restriction function of XPO1 is the delivery of nuclear RNAi and CIRF factors into VROs. These host restriction factors block the functions of viral RNAs and/or replication proteins, thus limiting the efficiency of replication. We show that the re-localization and antiviral functions of DRB4 and AGO2 RNAi components and CenH3 and nucleolin Nuc-L1 CIRFs depend on XPO1 nucleocytoplasmic function in *N. benthamiana* plants. These are selected proteins based on their known restriction functions [69, 70, 72, 73]. However, one can imagine that many more nuclear proteins, including those with antiviral or pro-viral functions, depend on XPO1 to be re-localized to the VROs. The emerging picture is that the co-opted XPO1 is a major protein interaction hub with a central role in deciding the outcome of tombusvirus infections by delivering a pool of restriction factors into VROs.

XPO1 was able to perform its antiviral function by binding to the viral p33 replication protein and using the co-opted actin network to target VROs. The recruitment of DRB4, AGO2 and CenH3 antiviral factors to VROs was dependent on both XPO1 and the co-opted functional actin filaments. The emerging theme on the roles of the co-opted actin network is that it facilitates the delivery of both cytosolic and nuclear proteins into VROs. These proteins provide either pro-viral or antiviral functions [75, 76]. Moreover, actin filaments also form “fences” within/around the vir-condensate and VROs, providing structural components to VROs [74]. Thus, the current work further expands our current understanding of the essential, but complex roles played by the co-opted actin network in tombusvirus replication.

Based on the recruitment of XPO1 and its antiviral cargos as well as XPO1 co-factors into tombusvirus VROs, we propose that XPO1 mobilizes the antiviral cargos from the nucleus to VROs (Fig. 13). We showed that the XPO1 co-factor Srm1/Prp20 guanine nucleotide exchange factor (RanGEF, known as RCC1 in humans) inhibits TBSV replication in yeast. RanGEF promotes the conversion of nuclear RanGDP into RanGTP, which binds to XPO1, resulting in conformational change by opening the cargo loading groove in XPO1. Then, the cargos are loaded via nuclear export signal (NES) sequences onto XPO1 [40, 43, 82]. The cargo-XPO1-RanGTP ternary protein complex is exported to the cytosol through nuclear pore complex. In the cytosol, the XPO1 complex takes advantage of the actin cytoskeleton, which is co-opted and stabilized by viral replication protein (TBSV p33 or CIRV p36) by inhibiting the depolymerization of actin filaments by COF1/ADF1 [63]. We propose that TBSV p33 interacts with XPO1 ternary complex in the cytosol and the complex is delivered to VROs along the co-opted actin filament networks. We showed that XPO1 regulatory cofactors, such as Ran GTPase-activating Protein1/2 (RanGAP1/2) and Ran Binding Protein1 (RanBP1), are also recruited into VROs. It is likely that the XPO1/cargo-complex dissembles in vir-condensate due to the stimulatory activities of co-opted RanBP1 and RanGAP1/2, releasing RanGDP and cargos from the XPO1 complex [40, 43]. FRAP analysis showed that the released antiviral cargo is present in vir-condensate substructure of the VROs. Therefore, the XPO1-delivered antiviral proteins could perform inhibitory functions within vir-condensate associated with VROs. In sum, we propose that XPO1 delivers and sequester antiviral cargos in vir-condensate (Fig. 13). Overall, the high concentration of antiviral cargos in vir-condensates enhances their effectiveness in inhibiting tombusvirus replication.

**Figure 13.**
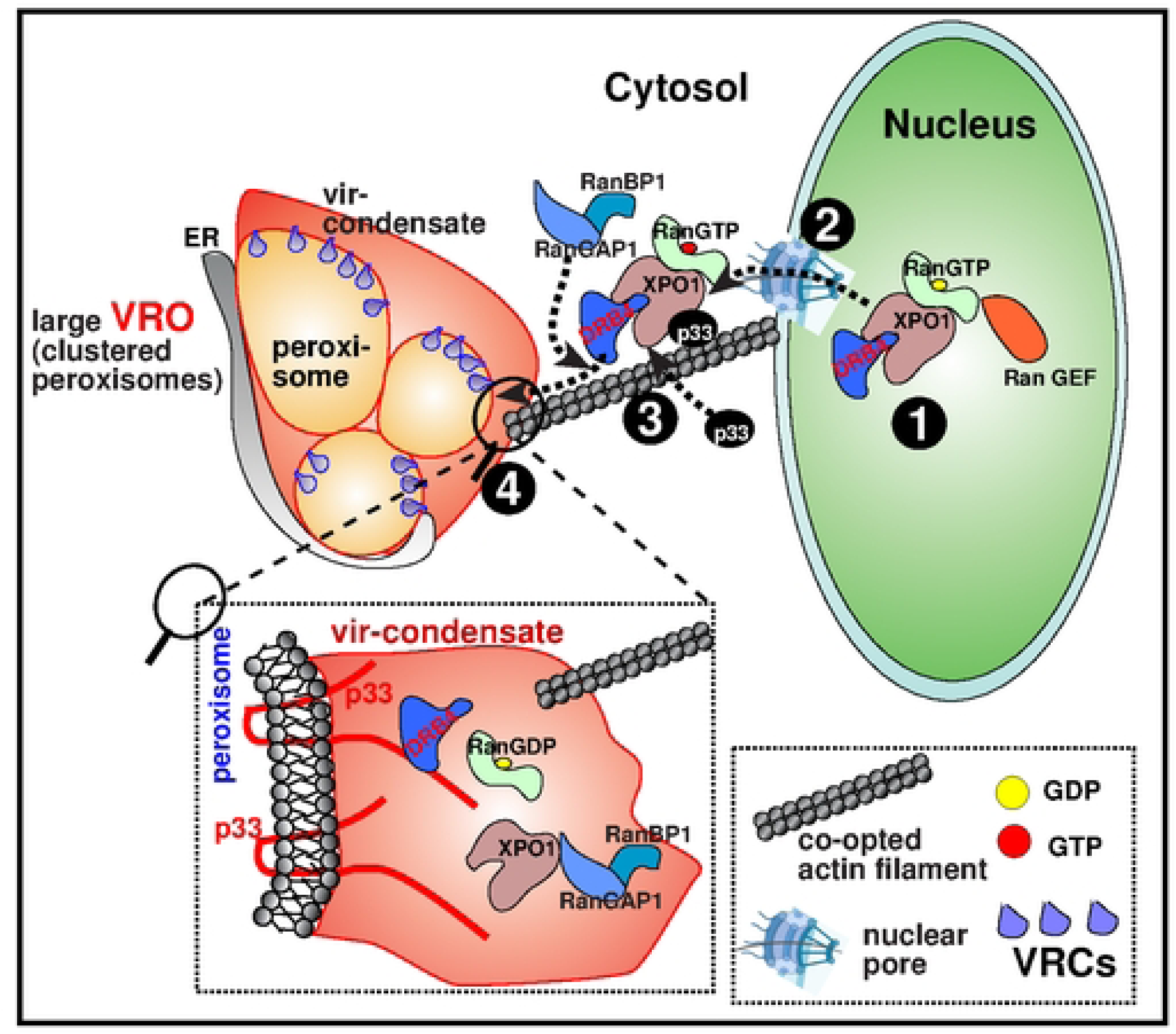
A model on the antiviral role of XPO1 during replication of tombusviruses. (#1) In the nucleus, high gradient of RanGTP is established with the contribution of guanine nucleotide exchange factor (RanGEF, also known as RCC1). RanGTP binding to XPO1 leads to conformational change opening the cargo loading groove in XPO1. Then, nuclear export signal (NES) sequence in the cargo is loaded onto XPO1, leading to the formation of cargo-XPO1-RanGTP ternary protein complex. (#2) Subsequently, this ternary protein complex docks at nuclear pore complex (NPC) in nuclear envelope, and then is exported to the cytosol through NPC. (#3) During replication of tombusviruses, the replication protein (TBSV p33 or CIRV p36) stabilizes the actin cytoskeleton by inhibiting the depolymerization of actin filaments by COF1/ADF1. TBSV p33 interacts with XPO1 ternary complex, resulting in recruitment of p33-cargo-XPO1-RanGTP to VROs along the stable actin filament networks. XPO1 regulatory cofactors, such as Ran GTPase-activating Protein1 (RanGAP1) and Ran Binding Protein1 (RanBP1), are also recruited into VROs represented by clustered peroxisomes (shown) or clustered mitochondria in case of CIRV infection (not shown). (#4 and the enlarged boxed area) The XPO1-complex dissembles in vir-condensate (induced by membrane-bound p33 or CIRV p36) due to the stimulatory activities of co-opted RanBP1 and RanGAP1, releasing RanGDP and cargos from the XPO1 complex. The released antiviral cargo performs inhibitory functions in vir-condensate associated with VROs. We propose that the host exploits XPO1 and the vir-condensate to concentrate and sequester antiviral cargos that enhances their effectiveness in inhibiting tombusvirus replication.

The established core roles of VROs are to support viral replication and protect the viral RNAs from degradation by host antiviral mechanisms. However, a major finding of this work is the demonstration of the abundant presence of antiviral factors within the vir-condensate substructure within the membranous VROs. Previous work demonstrated that vir-condensate contains recruited pro-viral cytosolic host factors, such as glycolytic and fermentation enzymes, which produce ATP locally to supply the energy requirement of tombusvirus replication [74]. The vir-condensate also stores autophagy proteins, such as NBR1 autophagy receptor and ATG8f core autophagy protein, in associated vir-NBR1 bodies to dampen the antiviral roles of autophagic degradation [83, 84]. However, based on the current work, the emerging picture on the role of the vir-condensate is becoming more complex: it seems to be a “central battleground” between the virus and the host for supremacy in controlling virus infection. We propose that the balance between the co-opted pro-viral and antiviral factors within the vir-condensate could be a major determining factor of virus replication and host susceptibility. We predict that by tilting the balance toward antiviral factors within vir-condensates might lead to new efficient virus therapeutics or host resistance against viruses.

## Summary

XPO1 exportin is a central interaction nod, which emerges as a major player in tombusvirus replication in plants. The emerging theme from our current studies is that XPO1 with the help of co-opted actin network provides nucleocytoplasmic transport of several viral restriction factors into the cytosolic VROs that restrict tombusviruses replication. The delivered restriction factors provide inhibitory functions within the vir-condensates associated with membranous VROs. Altogether, XPO1 is a critical protein interaction hub with major implications in viral replication.

## Materials and Methods

### Expression plasmids in yeast and plants

See details in the supplementary materials (S1 Text and S1-3 Tables).

### VIGS-based knockdown of *XPO1* in *N. benthamiana*

To silence expression of XPO1 in *N. benthamiana*, we used a virus-induced gene silencing (VIGS) approach [85, 86]. A 171-bp segment of *NbXPO1* (Accession number: LC434544) was cloned and inserted into pTRV2 vector to obtain the silencing vector pTRV2::NbXPO1 according to [87]. The leaves of *N. benthamiana* were infiltrated with a mixture of agrobacterium carrying pTRV1 (OD_600_ 0.2) and pTRV2::NbXPO1 (OD_600_ 0.2) or pTRV1 (OD_600_ 0.2) and pTRV2::cGFP (OD_600_ 0.2) as control for tombusvirus replication assay. Note that for the confocal microscopy experiments, in which NbXPO1 was knocked down in H2b-CFP or H2b-RFP transgenic *N. benthamiana*, the leaves were infiltrated with a mixture of agrobacterium carrying pTRV1 (OD_600_ 0.2) and pTRV::GST(OD_600_ 0.2) plasmids as the control. For virus replication assay, the upper leaves were inoculated with sap preparations containing TBSV, CIRV or CNV 7 d later. Note that we used CNV^20KSTOP^ not expressing the p20 suppressor of gene silencing protein [57]. The inoculated leaves were harvested at 1.5 dpi, 2 dpi and 2 dpi, respectively, to assay TBSV, CIRV and CNV RNA accumulation. For confocal laser microscopy analysis, the upper leaves were infiltrated with a mixture of agrobacterium expressing fluorescence protein fused to proteins of interest (OD_600_ 0.3) and mitochondrial or peroxisome markers (OD_600_ 0.3). The agroinfiltrated leaves were analyzed by confocal laser microscopy 2.5 days later.

### Transient expression of XPO1 to study replication of tombusviruses in *N. benthamiana*

To transiently express AtXPO1 (Accession number: AT5G17020) in the leaves of *N. benthamiana*, the coding region was cloned and inserted into pGD-eGFP-MCS vector to obtain pGD-eGFP-XPO1 (S1 Table). The leaves of *N. benthamiana* were infiltrated with a mixture of agrobacterium carrying pGD-eGFP-XPO1 (OD_600_ 0.5) and pGD-p19 (OD_600_ 0.2) to express AtXPO1. The mixture of agrobacterium carrying empty vector GD-eGFP-EV (OD_600_ 0.5) and pGD-p19 (OD_600_ 0.2) was infiltrated into the leaves as the control. To block the function of XPO1, the chemical inhibitor Leptomycin B (LMB) (Cell Signaling Technology, Inc.) was co-infiltrated with the agrobacterium mixture using the concentration of 400 nM, whereas 0.5% ethanol (EtOH) was co-infiltrated with the agrobacterium mixture as the control [88, 89]. The agroinfiltrated leaves were inoculated with sap containing TBSV, CIRV or CNV 18 h later. The samples were harvested at 1.5 dpi for TBSV and 2 dpi for CIRV and CNV, respectively, for total RNA or total protein extraction in the virus replication assay. Note that we applied a second, reinforcement infiltration of 400 nM LMB and 0.5% EtOH into the leaves 24 h after the first infiltration.

### Confocal microscopy imaging

To analyze the subcellular localization of XPO1 in healthy cells of H2b-CFP transgenic *N. benthamiana*, a mixture of agrobacterium carrying pGD-eGFP-XPO1 (OD_600_ 0.3), mitochondrial marker pGD-CoxIV-RFP (OD_600_ 0.3) or peroxisomal marker pGD-RFP-SKL (OD_600_ 0.3) and pGD-p19 (OD_600_ 0.2) were co-infiltrated into leaves. Subcellular localization of XPO1 during tombusvirus replication was done using leaves agroinfiltrated as above, followed by inoculation with sap containing TBSV, CNV or CIRV 12 h later. In protein co-localization experiments, we used a mixture of agrobacterium carrying pGD-p33-BFP or pGD-p36-BFP (OD_600_ 0.3), pGD-eGFP-XPO1 (OD_600_ 0.3), pGD-RFP-SKL or pGD-CoxIV-RFP (OD_600_ 0.2) and pGD-p19 (OD_600_ 0.2) for infiltration into the leaves of wild type *N. benthamiana*. The confocal microscopy imaging was performed at 2 or 2.5 post agrobacterium infiltration.

To analyze eGFP-XPO1 and its cargos in VROs during tombusvirus replication, a mixture of agrobacterium carrying pGD-eGFP-XPO1 or cargos (OD_600_ 0.3), pGD-RFP-SKL (OD_600_ 0.3), pGD-p19 (OD_600_ 0.2) together with 400 nM LMB or 0.5% (V/V) EtOH was infiltrated into the leaves of H2b-CFP transgenic *N. benthamiana*, followed by inoculation with TBSV or CIRV sap 12 h later. The confocal microscopy imaging was performed at 2 days post virus inoculation.

To analyze the subcellular locations of actin filaments and eGFP-XPO1 or its cargos during tombusvirus replication, the leaves of *N. benthamiana* were infiltrated with a mixture of agrobacterium carrying pGD-eGFP-XPO1 or cargos (OD_600_ 0.3), pGD-BFP-SKL (OD_600_ 0.3), LifeAct-RFP(OD_600_ 0.03) [90], and pGD-p19 (OD_600_ 0.2). The agroinfiltrated leaves were inoculated with TBSV 12 h later. Leaf samples were analyzed by confocal microscopy 2 days after TBSV inoculation. For CIRV infections, pGD-BFP-SKL (OD_600_ 0.3) was replaced by pGD-CoxIV-BFP.

To alter the actin filament network [91, 92], plasmids pGD-Flag-RavK (OD_600_=0.3) or pGD-Flag-VipA (OD_600_=0.3) carrying *Legionella* effector genes, either RavK or VipA, were co-infiltrated with the combinations mentioned above, followed by inoculation with TBSV or CIRV sap. The confocal imaging was performed 2 d later [93].

During the confocal microscopy analysis, CFP and BFP fusion proteins were excited at 458 nm or 405 nm, respectively, and detected at 460-490 nm or 430-470 nm, respectively. eGFP and RFP fusion proteins were excited at 488 and 561 nm and detected at a range of 495-535 nm and 560-600 nm, respectively, using an Olympus FV3000 microscope.

### BiFC (Bimolecular fluorescence complementation) assay

To test interaction between p33 and XPO1, a mixture of agrobacterium of pGD-nYFP-XPO1 (OD_600_ 0.3), pGD-p33-cYFP (OD_600_ 0.3), pGD-SKL-RFP (OD_600_ 0.3) and pGD-p19 (OD_600_ 0.1) was infiltrated into the leaves of *N. benthamiana,* followed by inoculation with TBSV sap 12 h later. To identify the interaction between p36 and XPO1, the leaves of *N. benthamiana* were co-infiltrated with agrobacterium carrying pGD-nYFP-XPO1 (OD_600_ 0.3), pGD-p36-cYFP (OD_600_ 0.3), pGD-CoxIV-RFP (OD_600_ 0.3), and pGD-p19 (OD_600_ 0.3), and were inoculated with CIRV sap 12 h later. The combination of pGD-nYFP-XPO1 and pGD-GST-cYFP was used as control. The leaf samples were analyzed with confocal laser microscopy at 2-2.5 dpi.

### Fluorescence recovery after photobleaching (FRAP)

Agroinfiltration of *N. benthamiana*. with various combinations of plasmids was done as described above. Note that the leaves were treated with 10 μM Latrunculin B (Abcam) to avoid the movement of VROs at least 3 h before microscopic imaging [94]. Photo-bleaching was performed in the middle area of VROs. Photobleaching process lasted ∼4-5 seconds with 405 nm laser at 80% intensity. And the confocal images were captured automatically by program for every 10 sec until 80 sec. For calculations, all the fluorescent intensity values were quantified using FLUO-view software installed on FV3000 Olympus confocal microscopy operation system. The calculation method for recovery rate was conducted according to [94]. The recovery curves including the recovery rate at different time points were plotted by GraphPad 9 software.

### Analysis of TBSV replication with *in vitro* reconstituted TBSV replicase in yeast cell-free extract (CFE)

Yeast cell-free extract (CFE) that supports in vitro TBSV repRNA replication was prepared with yeast strain BY4741 as described [95, 96]. Briefly, the *in vitro* reconstituted TBSV replicase assay was performed using a mixture of 2 μL of CFE, 0.5 μg DI-72 (+)repRNA, 0.2 μg affinity-purified maltose-binding protein (MBP)-p33 as well as MBP-p92^pol^ (both recombinant proteins were purified from E. coli), 5 μl of buffer A [30mM HEPES-KOH (pH 7.4), 150 mM potassium acetate, 5 mM magnesium acetate, 0.13 M sorbitol], 2 μl of 150 mM creatine phosphate, 0.2 μL of 10 mg/ml creatine kinase, 0.4 μl actinomycin D (5mg/ml), 0.2 μl of 1 M dithiothreitol (DTT), 0.2 μl of RNase inhibitor, 2 μl a ribonucleotide (rNTP) mixture (10 mM of ATP, CTP, and GTP as well as 0.25 mM UTP) and 0.1 μL of ^32^PUTP in a total of 20 μl reaction volume. The reaction was performed at 25°C for 3h and then stopped by the addition of 5 volumes of 1% SDS and 5 mM EDTA, followed by phenol-chloroform extraction and RNA precipitation. Then the repRNA products and dsRNA replication intermediates were analyzed by electrophoresis in 0.5X Tris-borate-EDTA (TBE) buffer in a 5% polyacrylamide gel (PAGE) containing 8 M urea.

## ACKNOWLEDGEMENTS

The authors thank Drs. W. Lin, and J. Pogany for valuable comments. We thank Dr. Boone (U. Toronto) for providing srm1 temperature-sensitive yeast mutant.

## Supplementary Materials

**S1 Fig. DRB4 and XPO1 are recruited to VROs during CNV replication.**

**S2 Fig. DRB4 and XPO1 are recruited to VROs during TBSV replication.**

**S3 Fig. DRB4 and XPO1 are recruited to VROs during CIRV replication.**

**S4 Fig. AGO2 and XPO1 are recruited to VROs during CNV replication.**

**S5 Fig. AGO2 and XPO1 are recruited to VROs during TBSV replication.**

**S6 Fig. AGO2 and XPO1 are recruited to VROs during CIRV replication.**

**S7 Fig. XPO1 delivers nuclear AGO2 to VROs to inhibit CNV replication.**

**S8 Fig. XPO1 delivers nuclear AGO2 to TBSV and CIRV VROs.**

**S9 Fig. CenH3 and XPO1 are recruited to VROs during TBSV replication.**

**S10 Fig. CenH3 and XPO1 are recruited to VROs during CNV replication.**

**S11 Fig. CenH3 and XPO1 are recruited to VROs during CIRV replication.**

**S12 Fig. XPO1 delivers nuclear CenH3 to VROs to inhibit CIRV replication.**

**S13 Fig. XPO1 is needed to deliver nuclear NUCLEOLIN to VROs to inhibit CNV replication.**

**S14 Fig. Association of DRB4 or AGO2 with co-opted actin filaments during tombusvirus replication.**

**S15 Fig. Association of CenH3 with co-opted actin filaments during tombusvirus replication.**

**S16 Fig. Actin filament network plays a critical role in delivering XPO1 or DRB4 antiviral cargo to CIRV VROs.**

**S17 Fig. Actin filament network plays a critical role in delivering CenH3 or AGO2 antiviral cargos to CIRV VROs.**

**S18 Fig. Regulatory co-factors of XPO1 are recruited into VROs during CIRV replication.**

**S19 Fig. Tombusvirus replication proteins do not interact with regulatory co-factors of XPO1 in *N. benthamiana*.**

**S20 Fig. XPO1 and the antiviral cargos are present in vir-condensates associated with membranous CIRV VROs.**

**S1 text.**

**S1 Table. List of plasmids constructed during this work.**

**S2 Table. List of plasmids from prior work.**

**S3 Table. List of primers used**.

